# Tissue-specific transcription footprinting using RNA PoI DamID (RAPID) in *C. elegans*

**DOI:** 10.1101/2020.08.19.257873

**Authors:** Georgina Gómez-Saldivar, Jaime Osuna-Luque, Jennifer I. Semple, Dominique A. Glauser, Sophie Jarriault, Peter Meister

**Affiliations:** University of Fribourg, Switzerland; Cell Fate and Nuclear Organization, Institute of Cell Biology, University of Bern, Switzerland; Institut de Génétique et de Biologie Moléculaire et Cellulaire (IGBMC); Institut National de la Santé et de la Recherche Médicale (INSERM) U1258/Centre National de la Recherche Scientifique (CNRS) UMR7104/Université de Strasbourg, France; Graduate School for Cellular and Biomedical Sciences, University of Bern, Switzerland

## Abstract

Differential gene expression across cell types underlies the development and cell physiology in multicellular organisms. *C. elegans* is a powerful, extensively used model to address these biological questions. A remaining bottleneck relates, however, to the difficulty to obtain comprehensive tissue-specific gene transcription data, since available methods are still challenging to execute and/or require large worm populations. Here, we introduce the <u>R</u>N<u>A</u> <u>P</u>ol Dam<u>ID</u> (RAPID) approach, in which the Dam methyltransferase is fused to a ubiquitous RNA polymerase subunit in order to create transcriptional footprints *via* methyl marks on the DNA of transcribed genes. To validate the method, we determined the polymerase footprints in whole animals, sorted embryonic blastomeres and in different tissues from intact young adults by driving Dam fusion expression tissue-specifically. We obtained meaningful transcriptional footprints in line with RNA-seq studies in whole animals or specific tissues. To challenge the sensitivity of RAPID and demonstrate its utility to determine novel tissue-specific transcriptional profiles, we determined the transcriptional footprints of the pair of XXX neuroendocrine cells, representing 0.2% of the somatic cell content of the animals. We identified 2362 candidate genes with putatively active transcription in XXX cells, among which the few known markers for these cells. Using transcriptional reporters for a subset of new hits, we confirmed that the majority of them were expressed in XXX and identified novel XXX-specific markers. Taken together, our work establishes RAPID as a valid method for the determination of polymerase footprints in specific tissues of *C. elegans* without the need for cell sorting or RNA tagging.

**Article summary:** Gene expression is a major determinant of cell fate and physiology, yet it is notoriously difficult to characterize in individual cell types for the widely used model system *C. elegans*. Here, we introduce a method based on the *in vivo* covalent modification of DNA by transcribing RNA polymerases to determine genome-wide transcription patterns in single tissues of embryos or young adult animals. We show that the method is able to identify actively transcribed genes in tissues representing down to 0.2% of the somatic cells in adult animals. Additionally, this method can be fully performed in a single laboratory by using third generation sequencing methods (ONT).

## Introduction

Differential gene expression across cell types encompasses both key determinants and markers of the cells’ identity. Cataloging these differences can provide critical insights and entry points in research aiming at elucidating the mechanisms controlling fundamental biological processes, such as organismal development and cell/tissue physiology. *C. elegans* is a widely used model animal, particularly well suited for integrative studies bridging our understanding across the molecular, cellular and organismal levels. With its transparent body, *C. elegans* was the first animal used to analyze tissue-specific transcription *in vivo* with GFP reporters (Chalfie *et al*. 1994), an approach still extensively used. In contrast to the versatility of the model for individual gene expression analysis, more holistic approaches such as tissue-specific transcriptomics still remain relatively challenging, in particular in postembryonic animals due to the tough cuticle and the difficulty to isolate intact tissue or cell types.

Two main general strategies have been developed to analyse specific tissues/cell types in *C. elegans*. A first general strategy involves the purification of tissue-specific messenger RNAs (mRNAs) from whole animals, which we will call “RNA tagging/pulling” hereafter. The most widely used RNA tagging/pulling method rely on the immunoprecipitation of a tagged poly-A binding protein-1 (FLAG::PAB-1) expressed in a specific cell type and cross-linked to RNA (Roy *et al*. 2002; Kunitomo *et al*. 2005; Pauli *et al*. 2006; Mattout *et al*. 2011; Hrach *et al*. 2020). One of the latest versions of this method is referred to as *polyA-tagging and sequencing* (PAT-seq; Blazie *et al*. 2015, 2017).The PAB-based method is technically demanding, in particular for the analysis of a small number of cells (Takayama *et al*. 2010) and has been shown to be associated with significant background noise (Von Stetina *et al*. 2007; Ma *et al*. 2016). Two more recent alternatives to PAB-based mRNA tagging are tissue-specific Translating Ribosome Affinity Purification (TRAP; Gracida and Calarco 2017; Rhoades *et al*. 2019) and trans-splicing-based RNA tagging (SRT; Ma *et al*. 2016). TRAP analysis focuses on ribosome-engaged mRNAs recovered after cross-linking and immunoprecipitation of a tagged ribosome subunit. TRAP allowed the identification of genes expressed in a specific neuron type representing only two cells per animal (Rhoades *et al*. 2019). SRT uses a modified SL1 splice leader expressed in target tissues which is trans-spliced to the transcripts by the cellular machinery (Ma *et al*. 2016). While SRT bypasses the noise inherent in immunoprecipitation procedures, it has so far only been applied to large tissues and the approach is limited to SL1-associated transcripts (62% of the C. elegans genes; Yang *et al*. 2017).

The second general strategy relies on the animal disruption and cell isolation, or dissociation, followed by the *in vitro* culturing or sorting of labelled cells (Von Stetina *et al*. 2007; Spencer *et al*. 2011), or nuclei (Haenni *et al*. 2012; Steiner *et al*. 2012) prior to transcriptomics analysis, and which we will call “dissociation-based” methods. The tough cuticle in larval stages and *a fortiori* in adults constitute a significant obstacle, and initial studies focused on embryonic cells which were more easily dissociated. More recent protocols combining FACS and RNA sequencing were successfully used to analyze major tissues (Kaletsky *et al*. 2018) as well as neuronal subsets in adults (Wang *et al*. 2015; down to 6 neurons per animal; Kaletsky *et al*. 2016). Combined with single-cell RNA sequencing (scRNA-seq), large-scale dissociation-based studies can address transcript profiles in multiple cell-types at the same time (Cao *et al*. 2017; Packer *et al*. 2019), including the analysis of individual neuron types (Hammarlund *et al*. 2018; Lorenzo *et al*. 2020). However, how efficiently specific cell types, especially rare cells, can be purified varies depending on their morphologies, how fragile they are and the developmental stage considered, suggesting that dissociation-independent methods could be useful complementary approaches.

We investigated whether we could use DNA adenine methylation identification (DamID) to footprint actively transcribed genes in specific cell types in whole worms. This approach has been successfully used in *Drosophila t*o determine transcribed genes in rare brain cell types (Southall *et al*. 2013). DamID relies on the low-level expression of a fusion between the *E. coli* Dam methyltransferase and a protein of interest, here a subunit of the RNA polymerase. Binding of the latter to DNA leads to the methylation of GATC sites in the vicinity of the binding site (Figure 1A). After DNA extraction, methylated GATCs are specifically cleaved by the restriction enzyme *Dpn*I and amplified using adapter-mediated PCR before sequencing (Figure S1A). The adaptation of the method to *C. elegans* seemed promising since endogenous adenine methylation/demethylation is very rare in worms and not targeted to GATC motifs (Greer *et al*. 2015). The GATC site frequency is moreover expected to provide good spatial resolution. Indeed, *C. elegans* has 269,049 GATC sequences per haploid genome, corresponding to an average of one site for every 374 bp and a median of 210 bp (Gómez-Saldivar *et al*. 2016). In *C. elegans*, DamID has been used so far to study the genomic footprint of the DAF-16 transcription factor (Schuster *et al*. 2010), and large-scale interactions between the genome and the nuclear periphery (Towbin *et al*. 2012; Sharma *et al*. 2014; Cabianca *et al*. 2019; Harr *et al*. 2020).

**Figure 1.**
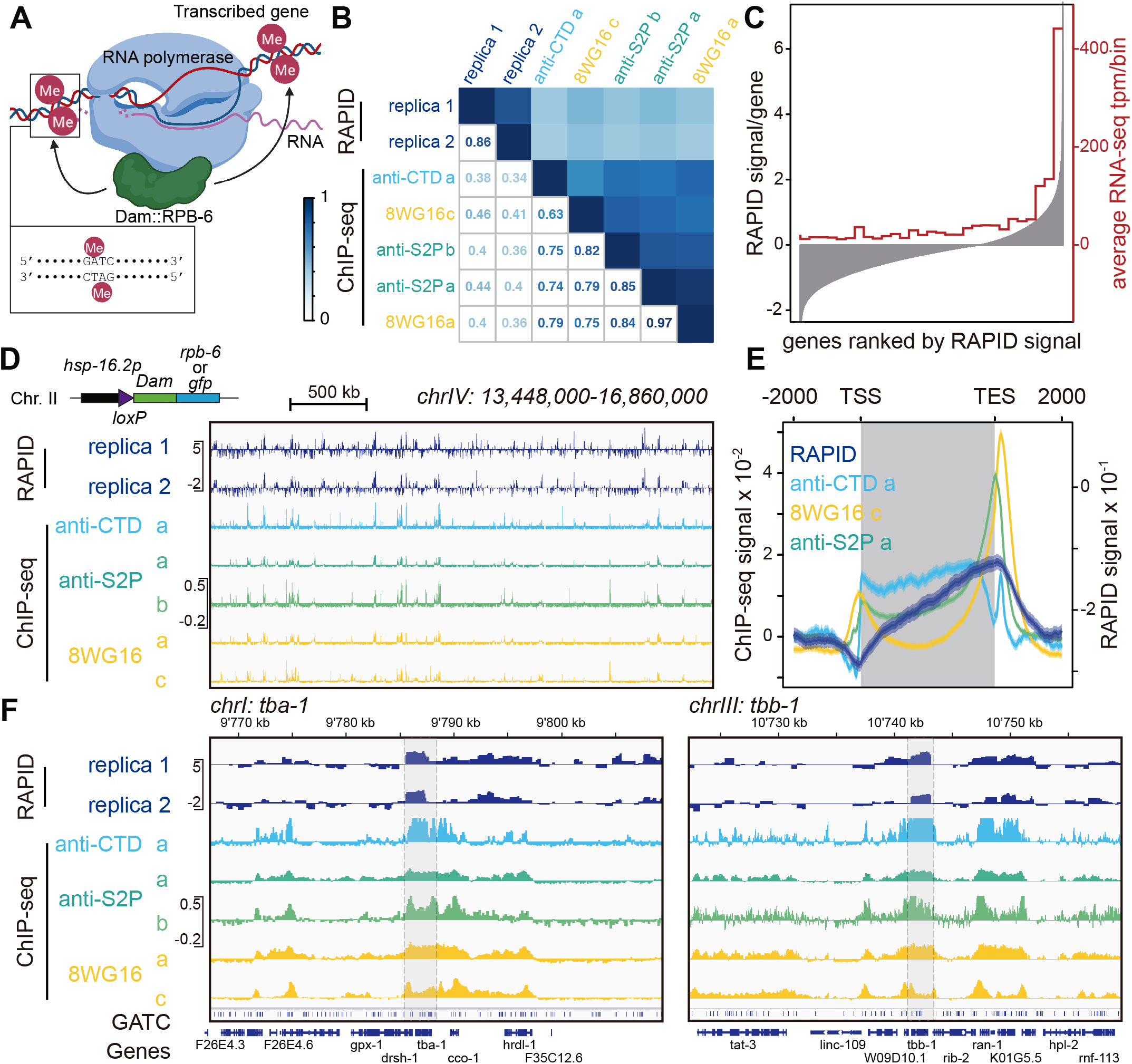
RAPID: <u>R</u>N<u>A</u> <u>P</u>oI II Dam<u>ID</u> scheme. **A.** *Dam::rpb-6* bound to all three RNA polymerase types modifies proximal GATC motifs. DamID analysis provides RNA polymerase footprints *in vivo*. **B.** Gene-level correlations in young adults whole animals of RAPID and RNA polymerase Chromatin immunoprecipitation-seq using different antibodies (a: (Garrido-Lecca *et al*. 2016); b: (Kalinava *et al*. 2017); c: (Miki *et al*. 2017)). **C.** Comparison of RAPID signal with mRNA expression profiling. All genes are represented on the x axis ranked, from left to right, based on RAPID signal in the entire animal (shown on left y axis). Averages transcripts per million (tpm, calculated with salmon) using dataset from study c above (rRNA depleted) were calculated, using the genes falling into each bin of 690 genes on the x axis (values on right y axis). **D. F.** Profiles at large scale (D) and gene scale (housekeeping genes, F) of RAPID and the different RNA polymerase II ChIP-seq studies cited in B. **E**. Metagene plot of the RNA polymerase II ChiP-seq and RAPID signals on WS270 genes. TSS: transcription start site; TES: transcription end site.

Here, we describe the RAPID approach (<u>R</u>N<u>A</u> <u>P</u>olymerase <u>D</u>am<u>ID</u>) for transcriptional footprinting in specific tissues in *C. elegans* in both embryonic blastomeres and in young adults. Using a fusion between a small subunit present in all three RNA polymerases, we show that the technique can be used on both fluorescently sorted blastomere cells, or DNA isolated from entire young adults using cell-type-specific expression generated by *Cre/lox*. To test the versatility of the method, we determined the polymerase footprints in three different tissues at each stage. In young adults, these tissues represent between 10 and 0.2% of the somatic cells of the animal. We show that meaningful transcriptional patterns can be recovered using this technique, in line with previously used, RNA tagging/pulling and dissociation-based methods. We further explore how this technique can be used to discover tissue-specific markers and report here the identification of eight new reporters for adult XXX cells. Additionally, as a result of the phasing-out of older sequencers able to sequence amplicons of different sizes as produced by DamID-seq, we show that new long read sequencing methods can be used to sequence DamID libraries, which makes it possible to carry out the experiments, from DNA extraction to amplicon sequencing in less than a week without any external sequencing facility.

## Materials and methods

### Plasmids and transgenic strains

All plasmids and worm strains are listed in Table S1 and S2. Tissue-specific DamID plasmids were generated using Gibson assembly. All DamID plasmids were integrated as single copies using MosSCI either on chromosome II or IV (see details in Supplementary Table 1; Frøkjaer-Jensen *et al*. 2008). Strains and plasmids are available upon request.

### Worm growth

Worms were grown on solid NGM, seeded with OP50 bacteria for maintenance culture and genetic crosses. All worm cultures were grown at 20°C. For cell sorting, animals were grown on peptone plates seeded with HB101, synchronized twice by hypochlorite treatment. Young adults containing up to 10 embryos were recovered 66 hours after plating synchronized L1 larvae. For DamID experiments in young adults, worms were grown on NGM seeded with Dam negative *E. coli* GM48 for at least two generations. Around 4’000 synchronized L1s were seeded onto 100 mm plates (1,000-1,200 per plate) and collected 53h later. Worms were washed extensively with M9 (at least 10 times) and distributed in aliquots of 30 μl removing the excess liquid. Samples were snap-frozen and stored at −80 °C.

### Sorting of embryonic blastomeres

Synchronized gravid adult hermaphrodites containing 8-10 eggs were treated with hypochlorite. Eggs were then incubated for 3 hours in M9 at 25°C until they reached the 1.5 fold stage. Eggs were transferred to 500 μl egg buffer (25mM Hepes pH7.3, 118mM NaCl 48mM, KCl 2mM, Cacl2 2mM MgCl2 in 250 ml) and pelleted 1 min at 2000 rpm. The supernatant was then aspirated, leaving 100 μl of buffer with the pellet. 500 U of chitinase (Sigma-Aldrich;C8241-25U) were added and the mixture resuspended and further incubated for one hour at room temperature. Chitinase was neutralized with 800 μl Leibovitz medium. Digested embryos were recovered by centrifugation at 3000 rpm for 5 minutes at 4°C. Embryos were then dissociated into isolated blastomeres by pipetting up and down with a P1000 pipette, up to 150 times until dissociation was complete. The cell population was filtered with a Millex-SV syringe 5 μm filter (Millipore;SLSV025LS). 500 fluorescent cells (one technical replicate) were sorted using a BD FACSAria Fusion (8000 events/second; 85μm nozzle) in sterile PCR tubes containing 1μl of pick buffer (50mM Tris-HCl pH 8.3; 75mM KCl; 3mM MgCl2; 137mM NaCl). Collected samples were then frozen in liquid nitrogen before further processing. After DamID processing and PCR (see below), two technical replicates were pooled for sequencing library preparation.

### DamID amplification, library preparation, and sequencing

For sorted cells, frozen samples were lysed by addition of 2 μl of lysis buffer (10mM TrisAc, 10 mM MgAc, 50 mM KAc, 0.67% Tween 20, 0.67% Igepal + 1mg/ml Proteinase K) and incubated for 2 hours at 60°C before Proteinase K inactivation at 95°C for 15 minutes. DamID amplicons were obtained as previously described (Gomez-Saldivar *et al*. 2016) using 30 PCR cycles.

For young adults, DamID was performed on 500 ng gDNA extracted from the animals using DNeasy Blood and Tissue Kit (Qiagen #69504). Two replicates for each stage and cell type were processed. DamID amplicons were obtained as previously described (Gomez-Saldivar *et al*. 2016) with 20 PCR cycles for worm-wide DamID, 22 PCR cycles for muscle DamID, 22-24 for intestine DamID and 24-26 cycles for DamID of XXX cells. New Illumina patterned flow cell technology does not allow sequencing of amplicons larger than 600 bp. Both types of DamID PCR amplicons were therefore sequenced using nanopore sequencing. After AMPure XP (Beckman-Coulter) purification with 1.8 bead volume, DamID PCR amplicons were directly used for nanopore library barcoding and preparation using NBD-104 and LSK-109 kits (Oxford Nanopore). Libraries were sequenced to obtain at least 1 million reads per library on MinION using R9.4.1 flow cells. We compared the performance of both sequencing approaches by sequencing the same libraries with both old paired-end HiSeq2500 Illumina flow cells and nanopore sequencers. Coverage obtained by both approaches were similar (see snapshot of mapped reads in Figure S1E). However, nanopore sequencing allowed the direct sequencing of longer amplicons (Figure S1F), most likely because Illumina cluster amplification does not work well on molecules longer than 800 bp, even on old generation (HiSeq2500) flow cells. We compared whole animal DamID sample libraries generated using previously described strains expressing either a free GFP-Dam fusion or a perinuclear lamin fusion (Sharma *et al*. 2014). Statistical comparison of both techniques at the single restriction fragment level gave a Pearson correlation coefficient between short PE and long reads between 0.84 and 0.86 (Figure S1G).

### Bioinformatic analysis

Nanopore sequences were basecalled and demultiplexed using guppy 3.6 with the high accuracy model before mapping to the ce11 genome using minimap2 (Li 2018). Reads were considered as DamID amplicons when both ends mapped +/- 8 bp from a genomic GATC motif. Filtered libraries were then used to call polymerase footprinting values using the damidseq_pipeline package (Marshall and Brand 2015) using bam files (parameters: --bamfiles --extend_reads=0). The damidseq pipeline normalizes for GATC accessibility using a free Dam (in our case Dam::GFP) profile. Per-gene values were extracted using polii.gene.call (https://owenjm.github.io/damidseq_pipeline/) using WS270 gene annotations. Correlations between libraries and methylation footprinting values were made using *ad hoc* R scripts available upon request. Normalized footprinting tracks were visualized using IGV. For the comparison with ChIP-seq tracks, fastq files were downloaded from GEO, mapped to ce11, normalized for sequencing depth with a pseudocount of 8 and log2 normalized to the input. Final figure construction was made using IGV and Adobe Illustrator.

### Array construction and microscopy

Reporter plasmids were assembled using the 3-fragment MultiSite Gateway system (ThermoFisher Scientific). Promoters of candidate genes were PCR amplified from N2 genomic DNA. Primer sequences and promoter lengths are described in Table S3. Candidate promoters were cloned into pDONR-P4-P1R vector (Invitrogen) by BP recombination, generating slot 1 Entry vectors (Table S2). Final expression plasmids were generated by LR recombinations between promoters (slot 1), mNeonGreen fluorescent ORF (slot 2; dg353; Hostettler *et al*. 2017), unc-54 3’ UTR (slot 3; pMH473, gift from Marc Hammarlund) and pDEST-R4-R3. The control localization/expression plasmid *[sdf-9p::NLS::wrmScarlet]* was generated through an LR recombination reaction between dg801 (slot 1), dg651 (slot 2; Marques *et al*. 2019), pMH473 (slot 3), and pDEST-R4-R3. Finally, *promoter::mNeonGreen* and *sdf-9p::NLS::wrmScarlet* plasmids were co-injected into N2 young adult hermaphrodites at 20 ng/μl each, with 20 ng/μl of dg9 (*unc-122p::RFP*; red coelomocyte) as a co-injection marker (Addgene #8938; Miyabayashi *et al*. 1999). Two stable lines of each candidate promoter were selected (Table S1). For imaging, young adult animals identified based on the red coelomocyte reporter expression were transferred to a 2% agarose pad containing 0.01% sodium azide. Worms were then imaged on a Zeiss Axiovert microscope with a 20x objective driven by Visiview, using a Photometrics Coolsnap Myo CCD camera with GFP and RFP settings (60 planes spaced by 2 μm), plus a middle plane section in DIC settings. Using Fiji, the acquired optical stacks were then partially z-projected to capture the worm section containing the XXX cells, identified using the *sdf-9p::NLS::wrmScarlet* signal. Final figure construction was made using Adobe Illustrator. For each promoter except *asp-9*, two independent extrachromosomal arrays were scored in at least 5 animals.

### Identification of detected and unique tissue-specific genes and validation

Gene coverage of RAPID profiles are listed in Table S4 (blastomeres) and 5 (young adults) (Sheet 1-2). Detected genes in a tissue were defined as genes that are significantly expressed (FDR < 0.05) in one replicate of one tissue (Table S4, sheet 3 and Table S5, sheet 3). Genes consistently detected in a tissue were defined as genes that are significantly expressed (FDR < 0.05) in both replicates of one tissue (Table S4, sheet 4 and Table S5, sheet 4). Representative examples of consistently detected genes used in Figure 2C were selected from the CeNGEN project (https://cengen.shinyapps.io/SCeNGEA/) (Hammarlund *et al*. 2018) and visualized using IGV (https://igv.org/).

**Figure 2.**
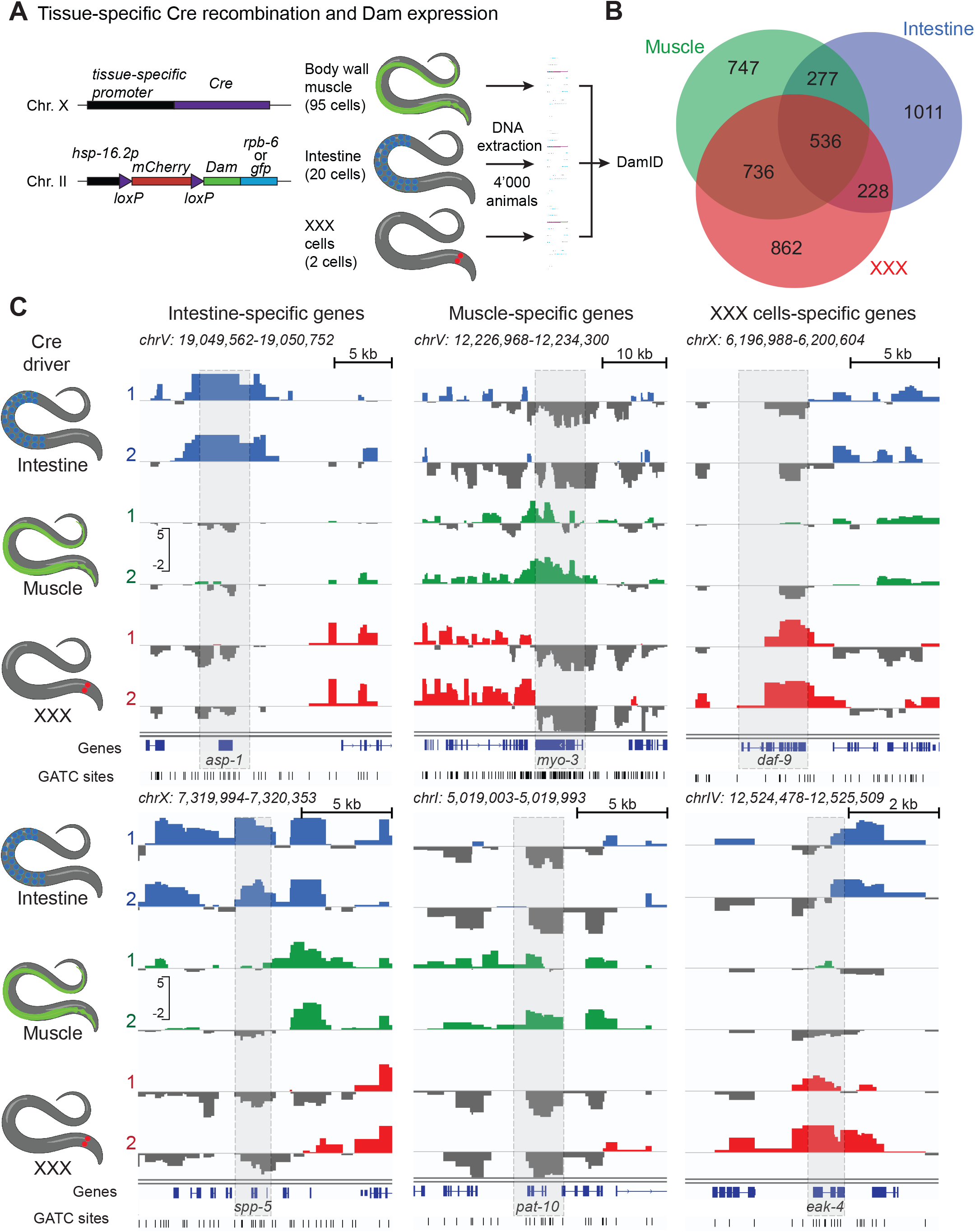
Tissue-specific expression profiles using RAPID by Cre/*lox* recombination in young adult animals. **A.** Experimental system fortissue-specific expression of Dam fusions and RAPID analysis. **B.** Venn diagram of overlap between expressed genes identified using RAPID in 3 different tissues. **C.** RAPID profiles for previously characterized genes expressed in a tissue-specific manner in intestine, muscle, and the XXX neuroendocrine pair.

Unique tissue-specific genes were defined as genes significantly and differentially expressed relative to the other two tissues (Table S4, sheet 5 and Table S5, sheet 5). Analysis of the uniqueness of the genes from each tissue was performed using the JavaScript library jvenn (http://bioinfo.genotoul.fr/jvenn).

To ensure an unbiased comparison between genes identified as expressed in muscle with RAPID and the set of genes identified with SRT (trans-Splicing Rna Tagging; Ma *et al*. 2016), only SL1 trans-spliced genes were selected, based on the intersection with the 10,589 mRNAs annotated as SL1-trans-spliced by modENCODE (Allen *et al*. 2011), generating a subset of 3477 genes. For the comparison between the genes identified in the intestine with RAPID and the set of genes identified with FANS (Fluorescence Activated Nuclear Sorting; Haenni *et al*. 2012), downregulated genes were eliminated from the list and only genes found as expressed and upregulated, in total 9169 genes, were analysed. For the comparison between the genes identified with RAPID and the set of genes identified with FACS (Kaletsky *et al*. 2018) in muscle and intestine, we directly used the expressed genes list from Table S1, which contained 7690 and 9603 genes, respectively. Chi-squared (χ2) statistical tests were performed using the GraphPad QuickCalcs Web site https://www.graphpad.com/quickcalcs/contingency1/ (accessed August, 2020).

### Tissue-specific prediction analysis

Tissue expression prediction analyses were performed using the top 500 statistically significant genes that were uniquely enriched in muscle and intestine (Table S5, sheet 6), selecting a multi-gene search within the Tissue-specific Expression Predictions for *C. elegans* program, version 1.0 (http://worm-tissue.princeton.edu/search/multi). For the tissue expression prediction test using the RAPID muscle-SL1 group, the parameters published in (Ma *et al*. 2016) were used.

### Gene ontology analysis

Gene Ontology analyses were performed on unique tissue-specific gene lists. GO terms and q-values were obtained using gProfiler (https://biit.cs.ut.ee/gprofiler/gost); (version e100_eg47_p14_7733820) with g:SCS multiple testing correction method applying a significance threshold of 0.05 (Table S5, sheet 7; Raudvere *et al*. 2019).

Functional enrichment analysis for genes detected with different methods was performed using WormCat (http://www.wormcat.com/index). Annotations classified into Category 2 were visualized with heatmap diagrams.

### Candidate selection for reporter analysis

Candidate genes were selected for follow-up promoter analyses with two goals in mind: demonstrating that they are indeed transcribed in XXX cells and identifying markers with expression restricted to XXX cells. Since XXX cells represent only 0.2% of the *C. elegans* cellular content, we reasoned that XXX-specific markers should produce very little or no signal in the whole animal samples. Therefore, out of the 862 XXX-enriched genes, with a significant RAPID signal in XXX, but not in muscle or in intestine samples, we further removed genes detected in at least one worm-wide sample. We obtained a refined list of 275 XXX-marker gene candidates, which still included the known XXX cell markers *daf-9* and *eak-4* (Table S5, sheet 8). From this candidate list, we selected *dhs-17* as an uncharacterized gene with a plausible link to XXX cell function, plus 11 random candidates. In order to limit caveats related to operons, we excluded a candidate if another transcript was located less than 200 bp upstream. The XXX-specific RAPID signals (RNA pol occupancy values) in the 12-gene subset (average=1.35, SD=0.59) were similar to those in the starting 275-gene set (average=1.47, SD=0.64; *p*=.16 by Student’s *t*-test).

### Data availability

Embryonic and adult DamID-seq data is available under the GEO accession GSE157418.

## Results

### RNA polymerase footprinting using RAPID

The RAPID approach relies on the expression of trace levels of the *E. coli* DNA adenine methyltransferase (Dam) fused to a subunit of the RNA polymerases, and the analysis of its DNA occupancy to evaluate the transcriptional state of genes. We first tested whether we could use AMA-1, the largest, catalytic subunit of RNA polymerase II, as reported in Targeted DamID (TaDa), done in neuronal lineages in *Drosophila* (Southall *et al*. 2013). However, the Dam signal was too weak, possibly due to the fact that the expression level of this very large fusion protein under transcriptional control of the uninduced *hsp-16.2* promoter was too low (not shown), or that the localization of the Dam domain within the *C. elegans* polymerase complex was not favorable to access its DNA substrate. We therefore replaced AMA-1 with RPB-6, the *C. elegans* homolog of the RNA polymerase subunit F present in all 3 RNA polymerases (personal communication M. Barkoulas; Katsanos et al., in preparation). RPB-6 together with RPB-1 and RPB-2, forms a ‘clamp’ that retains DNA near the active center of PolII (Cramer *et al*. 2000), stabilizing the transcription on the DNA template. In contrast to AMA-1 attempts, we obtained a strong DamID amplicon PCR signal in animals carrying a single copy transgene of the *rpb-6::Dam* fusion. As these animals had not been subjected to heat shock, they therefore expressed only low levels of the Dam fusion, yet the methylation levels were sufficient to perform DamID. Amplicons were sequenced using an ONT MinION nanopore device (see methods section for details and experimental validation of the technique). After signal normalization with GFP::Dam data to control for overall chromatin accessibility, we could detect general patterns of methylation that were consistent with polymerase-dependent methylation (Figure 1D-F) and a good correlation at the gene level between replicates (Figure 1B,D,F). We compared the RAPID profiles with published recent RNA polymerase II ChIP-seq datasets in young adult animals (Garrido-Lecca *et al*. 2016; Kalinava *et al*. 2017; Miki *et al*. 2017). Visually, the patterns appeared similar (see examples of profiles in Figure 1D,F). Even if the nature of the data is not identical, we performed a comparative analysis between the datasets for all genes, downscaling the resolution of the ChIP-seq data to GATC fragment (Figure 1B). RAPID and RNA polymerase ChIP-seq show a significant correlation (*R*=0.38-0.46). Overall, inter-replicate correlations were higher between the RAPID replicates (R=0.86) and between the RNA polymerase II ChIP experiments (R=0.63-0.97). The difference between the RAPID and RNA Pol II ChIP profiles most likely results from the different approach to recover DNA and from the fact that RAPID in adults represent a picture of cumulative transcriptional activity, while the RNA pol II ChIP represents a polymerase occupancy snapshot at the young adult stage.

To get a more quantitative understanding of the relation between RAPID and transcript abundance, we compared RAPID with RNA-seq data, using one of the published young adult RNA-seq datasets (Miki *et al*. 2017). We ranked all genes based on their RAPID signal and calculated the average RNA-seq signal across 30 equally-sized bins. We observe a clear correlation between RAPID and RNA-seq: genes with high RAPID signal are highly expressed as determined by RNA-seq, while lowly-expressed genes harbor low RAPID levels (Figure 1C).

To understand how the methylation signal spreads along the body of the genes, we constructed metagene plots over all 20’000 *C. elegans* genes for RAPID and RNA pol II ChIP-seq experiments. ChIP-seq signals show a characteristic profile, as the polymerase accumulates at the transcription start sites and at the transcription end sites (TSS and TES, respectively, Figure 1E). RAPID signal increases steadily from the TSS to the TES where it peaks before decreasing from the TES onwards (Figure 1E). This 3’ end accumulation is similar to the one observed in ChIP-seq. In contrast, the difference between ChIP-seq and RAPID profiles (absence of a 5’ peak in RAPID) is very likely due to the localization of the RPB-6::Dam fusion inside the RNA polymerase complex. Located on the opposite side of the RPB-1/2 complex relative to the DNA strands, the presence of the pre-initiation complex (PIC) at the promoter greatly restricts access to DNA (for review: Cramer 2004; Schier and Taatjes 2020). Once the polymerase switches to elongation, leaving the PIC on the promoter, Dam is likely to gain access to the DNA and efficiently methylates the transcribed region, as observed in the RAPID profiles.

We also examined the profile in genes transcribed by Pol I and Pol III. As expected from the differential inclusion of AMA-1 and RPB-6 subunits in the different polymerases, RAPID with RPB-6 also labelled genes transcribed by RNA Pol I and III. RAPID showed high enrichments on the rRNA genes at the end of chromosome I (Figure S2A), as well as a majority of previously characterized snoRNAs transcribed by RNA polymerase III (46 out of 57; Ikegami and Lieb 2013). snoRNA are sometimes located in introns of Pol II-transcribed genes, making it difficult to differentiate whether the RAPID signal originates from the snoRNA transcription or from the overlapping gene. Nevertheless, 24 out of 44 RAPID-positive snoRNA genes could be unambiguously assigned to Pol III transcription as there was no overlap (Figure S2B). In contrast, the corresponding signal was markedly weaker in the young adult RNA pol II ChIP-seq datasets and very high for the RNA Pol III subunit RPC-1 (performed in embryos, Figure S2B; Ikegami and Lieb 2013).

Taken together, these data show that the RAPID method is suitable to reveal RNA polymerase footprints, serving as indirect indications of transcriptional activity by the three different RNA polymerases.

### RAPID in FACS-sorted embryonic cells

Since RAPID could label transcribed genes in whole animals, our next goal was to adapt the method for tissue-specific analysis. We first explored whether RAPID could be used to footprint transcription in specific cell types of the embryo purified using fluorescent sorting. Using strains constitutively expressing Dam fusions, we sorted 1000 cells for 3 different tissues of various abundance: body wall muscle, intestine and the rectal Y cell (80, 20 and 1 cell per embryo, respectively; Figure S3A, Table S4). RAPID could produce footprints for all samples (Figure S3C), and the reproducibility of the RAPID signal at the gene level between replicas was good for the intestinal sample (*r=0.78*) yet lower for muscle and Y cell samples (*r=0.46* and *0.24*, respectively; Figure S4A). When using an FDR below 0.05, we detected a total of 4986, 3165 and 4819 genes with a high RAPID signal in the muscle, intestinal and Y datasets respectively, and 1566, 1570 and 840 consistently detected genes (FDR<0,05, in both replicates) in these tissues (Table S4). In these experimental conditions, RAPID appears suitable to identify transcription footprints of cell type specific genes, such as the intestinal genes *cpr-6* (cathepsin B; Pauli *et al*. 2006) and *elo-2* (fatty acid elongase ELOVL; Han *et al*. 2017), or the Glutathione S-Transferase gene *gst-4* expressed in muscle, as well as the muscle-specific WDR1 homolog *unc-78* (Figure S3C; Mohri and Ono 2003; Hasegawa *et al*. 2007). Comparison of the footprints obtained for all three tissues allowed us to define 395 Y-specific genes (Figure S3B), and among them, shallow Y-specific RAPID signal was visible for *ceh-6* and *sox-2*, two transcription factors previously shown to be involved in Y-to-PDA transdifferentiation initiation (Figure S3C; Kagias *et al*. 2012). In contrast, we observed some unexpected RAPID footprints for genes which we expected to be expressed in a tissue-specific manner. For example, the myosin *myo-3* was observed in all 3 tissues (Figure S3D). Conversely, some genes which are known to be expressed in tissue-specific manner did not show high RAPID values, *e.g*. the intestine-specific *asp-1* and *spp-5* genes or the muscle-specific *pat-10* one (Figure S3D). These differences might have a biological significance: most transcription patterns have been defined in older animals, yet those genes might be expressed more broadly, or not at all, in embryonic blastomeres. Alternatively, the lack of a robustly established transcription pattern might hinder faithful footprinting. Indeed, RAPID requires the enzymatic action of the methyltransferase to build up the signal, while footprints are removed at each replication round in rapidly dividing blastomeres. Together with the stochasticity of single methylation events, this might decrease the precision of the profiles. In agreement with this, intestinal cells, most of which are born and remain postmitotic early, display high reproducibility in the RAPID signal, while late-born muscle and the Y cell are less reproducible (Figure S3C,D). In conclusion, RAPID allows transcriptional footprinting of sorted cells, provided cells have been born long enough for the Dam fusions to methylate reproducibly their target genes. These results thus suggest that RAPID may be more suited for the study of later developmental stages.

### Tissue-specific RAPID in young adult animals

We next asked whether we could use the RAPID approach for the analysis of tissue-specific transcription profiles in adult worms. To circumvent the technical difficulties associated with mature animal dissociation and cell sorting, we sought to implement a transgenic approach for the cell-specific targeting of the RPB-6::Dam fusion. Several strategies have been developed to target Dam fusion expression to specific tissues, using either bicistronic constructs attenuating Dam translation (Southall *et al*. 2013), tamoxifen-induced nuclear translocation (Pindyurin *et al*. 2016; Redolfi *et al*. 2019) or tissue-specific recombination cassettes (Muñoz-Jiménez *et al*. 2017). We used the latter strategy using an insertion of a floxed mCherry ORF located between the *hsp-16.2* promoter and the Dam fusion (Figure 2A). Tissue-specific expression of the Cre recombinase leads to low-level expression of the Dam fusion (Ruijtenberg and van den Heuvel 2015). We used three different Cre drivers expressed in muscle, intestine and a pair of head neuroendocrine cells called XXX, using *myo-3*, *elt-2* and *sdf-9* promoters, respectively. We conducted cell-specific RAPID with only 4’000 young adult hermaphrodites as starting material for DNA extraction. DamID signal across genes was highly correlated between replicates, and less correlated between different Cre drivers, suggesting tissue-specific DamID footprinting (Figure S4B).

Among the genes with high RAPID signal, we defined sets of *detected* genes in each tissue (with a FDR below 0.05 in at least one replicate), representing 4379, 4583 and 3901 genes with high RAPID footprint in intestine, muscle and XXX cells, respectively. Among these, we also defined sets of *consistently detected* genes (with FDR of <0.05 in both replicates), encompassing 2052, 2296 and 2362 genes in intestine, muscle and XXX cells, respectively. Out of these, 1011, 747 and 862 were detected in only one tissue and defined as *tissue-specific* (Figure 2B, Table S5, sheet 5). 536 genes harbored high RAPID footprints in all three sampled tissues (Figure 2B, Table S5, sheet 9), and a GO term analysis highlighted many very general terms consistent with the function of housekeeping genes (Table S5).

We validated these results focusing first on individual genes with well-characterized tissue-specific expression (Figure 2C). The intestine-specific prograstricsin aspartyl protease homolog gene *asp-1* and the saposin-like channel defense gene *spp-5* were exclusively footprinted in intestinal samples (Tcherepanova *et al*. 2000; Roeder *et al*. 2010). *myo-3*, the muscle myosin gene and *pat-10*, a troponin gene, both exclusively expressed in muscle tissues, harbored high RAPID footprints in muscle samples (Miller *et al*. 1986; Kagawa *et al*. 1997). Finally, the XXX-specific genes *daf-9* (encoding a P450 cytochrome homolog) and *eak-4* (encoding a membrane protein) were marked with RAPID exclusively in XXX cells (Gerisch *et al*. 2001; Hu *et al*. 2006).

### Comparison of tissue-specific RAPID with available datasets in muscle and intestine

To expand the validation of our RAPID footprint gene lists beyond selected tissue markers, we next engaged three more comprehensive approaches. These approaches focused on muscle and intestine, since no large-scale gene expression data are available in XXX cells. First, we focused on a narrower set of genes with high RAPID footprint present exclusively in one of the tissues. These gene lists with the top 500 unique genes of each tissue, were analyzed with a computational method that predicts tissue-specific enrichment, using a multi-gene query (http://worm-tissue.princeton.edu/; Chikina *et al*. 2009). The tissue prediction tool showed that muscle-only and intestine-only RAPID footprinted genes gave a high prediction score for muscle and intestine, respectively (Figure 3A, B). In contrast, their scores were much lower for non-targeted tissues (including neurons, hypodermis and germline).

**Figure 3.**
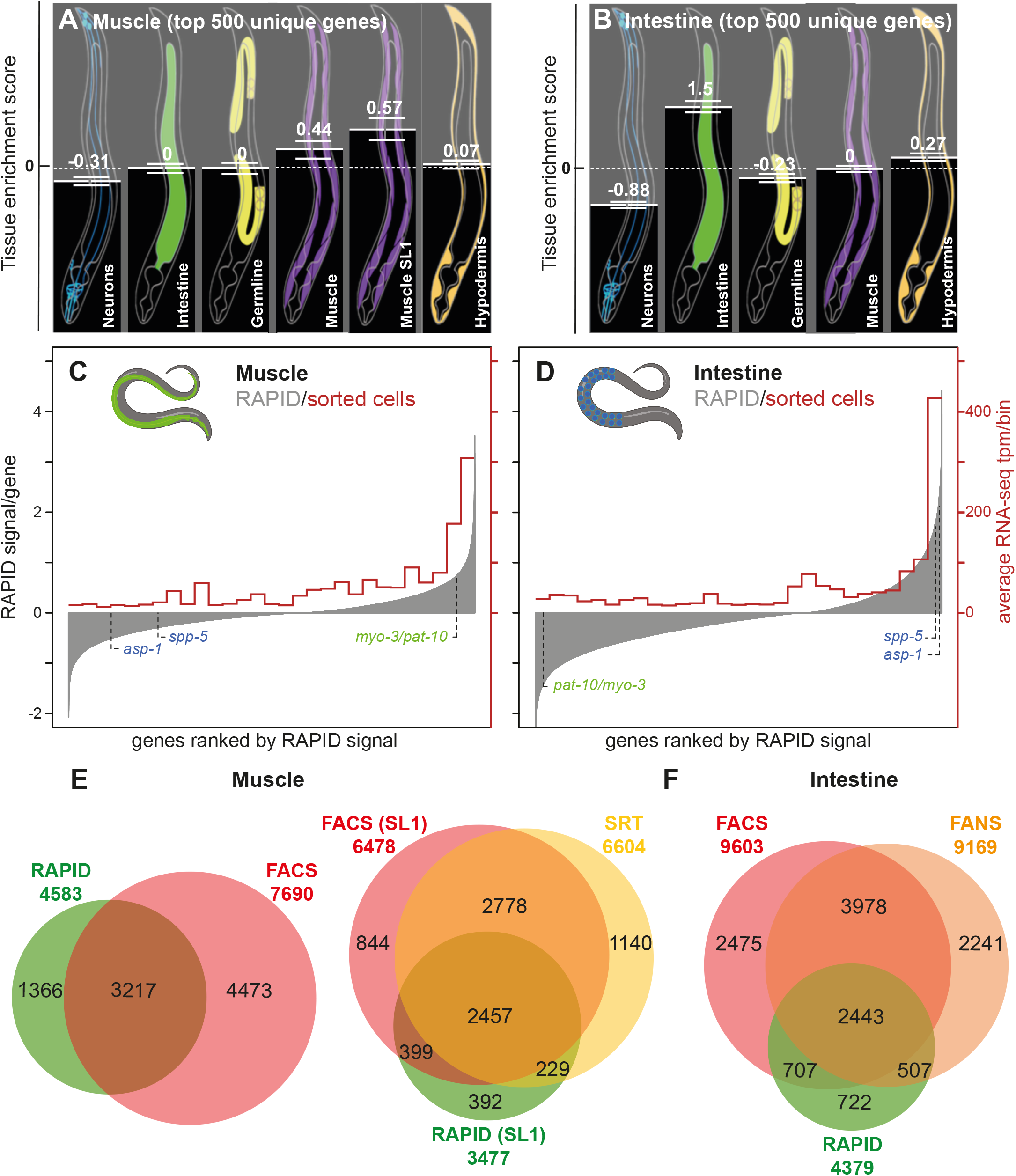
Validation of RAPID muscular and intestinal transcriptomes. **A-B**. Tissue enrichment prediction of tissue-specific RAPID transcriptomes. The top 500 hits of unique tissue-enriched genes for muscle (A) and intestine (B) were analyzed by the online tool Tissue-specific Expression Predictions for *C. elegans* (http://worm-tissue.princeton.edu/search). Tissue enrichment scores (mean and SEM) showed that the top genes of each tissue identify their corresponding original tissue. **C-D.** Semi-quantitative comparison between RAPID and FACS/RNA-seq in intestine and muscle (Kaletsky *et al*. 2018). All genes are represented on the x axis ranked, from left to right, based on the RAPID signal in the considered tissue (shown on left y axis). Averages of transcripts per million (tpm) for those genes were calculated, binning genes by pools of 690 (30 bins) on the x axis (values on right y axis). The muscular and intestinal marker genes presented in Fig. 2C are indicated in green and blue, respectively. **E.** Overlap of genes footprinted using RAPID with muscle-expressed genes detected by FACS (left; (Kaletsky *et al*. 2018) and SRT (right; Ma et al. 2016). For the comparison to SRT data, the subsets of SL1-trans-spliced genes detected using RAPID and FACS were used. **F.** Overlap of genes footprinted using RAPID with intestine-expressed genes detected by FACS (Kaletsky et al. 2018) and FANS (Haenni *et al*. 2012).

Second, we compared the genome-wide RAPID footprint signal levels with expression levels of genes determined by RNA-seq in FACS-sorted body wall muscle and intestinal cells (Kaletsky *et al*. 2018). As for whole animal samples (Figure 1C), we observed a correlation between RNA-seq levels and RAPID signals for both tissues (Figure 3C,D). Similar correlations were observed when comparing RAPID signal with single cell RNA-seq data (scRNA-seq, Figure S8), with the caveat that the scRNA-seq experiment was performed in the second larval stage, not in young adults (Cao *et al*. 2017).

Third, we compared the RAPID-detected gene sets with those reported for adult muscle and intestine in RNA-seq-based studies with (RNA-seq of FACS-sorted cells by Kaletsky *et al*. 2018), SRT (Ma et al., 2016), FANS (Haenni et al., 2012) or PAT-seq (Blazie *et al*. 2017). A majority of genes with detectable RAPID footprints were also identified with these tissue-specific transcriptomic methods. We found that 90% of the muscle RAPID hits (Figure 3E and Figure S5), and 87% of the intestinal RAPID hits were detected by at least one of the other methods (Figure 3F and Figure S5). Remarkably, the overlap between the genes detected by RAPID and other methods is in the same range as that found between other methods (our own data in Table S5, and similar analyses in Haenni *et al*. 2012; Ma *et al*. 2016). Although these results are encouraging and suggest that RAPID hits are genuinely expressed in these tissues, we note that the number of genes detected with RAPID is overall lower than that detected with other methods, especially in the intestine, where FACS-seq and PAT-seq detect twice as many genes as RAPID (Figure S5C). To evaluate what kind of genes were not identified by RAPID under our experimental conditions, we performed GO-analyses on the common gene sets (identified by two methods) and method-specific gene sets (identified by one but not the other method) (Figure S6 and S7, and Table S6). For this purpose, we used the WormCat analysis tool (Holdorf *et al*. 2020), specifically developed to categorize and visualize *C. elegans* GO data. For the RAPID/FACS and RAPID/PAT-seq common gene sets, we mostly found GO categories related to widely-expressed genes (*Development, Transmembrane proteins, Signaling, Metabolism*…) and categories that one would expect in these tissues (Figure S6/S7). For example, significantly enriched categories in muscle included: *Muscle function* and *Cytoskeleton*; and in the intestine: *Metabolism of lipids*, *Trafficking of vesicles*, *Endocytosis*, and *Lysosomes* (Figure S7). GO categories associated with RAPID-missed genes (only-FACS, only-PAT-seq in Figure S7) and FACS/PAT-seq common genes largely overlapped with those from the RAPID/FACS and RAPID/PAT-seq common genes, and contained additional very general categories (such as *Transcription, DNA, Ribosome, Development* or *mRNA processing*). They also included less expected categories (such as *Extracellular material: collagen, Neuronal function: neuropeptide* and *Major sperm protein* in muscle, and *Muscle function, Cytoskeleton actin function*, and *Major sperm protein* in intestine). These latter categories may point to possible contaminations by other tissues, a known phenomenon in cell-sorting and cross-linking-based methods (Von Stetina *et al*. 2007; Haenni *et al*. 2012; Spencer *et al*. 2014).

Taken together, the results of our comparative analyses with previous gene expression studies indicate that, even if the number of RAPID-detected genes tends to be lower than for RNA-seq-based methods, RAPID identifies tissue-specific RNA polymerase transcriptional activity that largely matches the specific transcriptional output in these tissues.

### RAPID identifies genes expressed in XXX cells

XXX cells are neuroendocrine cells derived from hypodermal embryonic progenitors. They express enzymes required for steroid hormone synthesis such as DAF-9 and are implicated in the control of dauer formation during development (Gerisch *et al*. 2001; Ohkura *et al*. 2003). As we still have limited information on the set of genes expressed in XXX cells, as well as on the role of those cells in adults, our XXX-cell transcribed gene dataset could represent a valuable resource for future studies. We thus conducted additional experiments to confirm the validity of RAPID profiles to infer gene transcription in those cells and identify new markers.

A GO term analysis highlighted many terms related to secretory and neural functions in the gene set specific for XXX cells, while they were absent from the muscle- and intestine-enriched gene sets (Table S5, sheet 7). These results are expected for neuroendocrine cells, with functional properties very similar to those of neurons.

To further confirm the validity of RAPID profiles to infer gene transcription, we selected 12 genes with high RAPID signal in XXX cells compared to intestine, muscle and whole animals, by comparing their profiles in the four tissues (Figure S6). We cloned their putative promoters in front of a mNeonGreen fluorescent reporter (Hostettler *et al*. 2017) and created transgenic lines bearing extrachromosomal arrays by gonad microinjection, together with an XXX-specific red reporter driven by the *sdf-9* promoter (the promoter which we previously used for XXX-specific RAPID). Out of the 12 promoters, eight drove expression in the XXX cells at various levels (*asic-1, dhs-17, F14H8.2, C06A1.3, asp-9, B0034.5, F52E1.5*, and *nlp-15*, Figure 4), one was impossible to score (*T22E5.6*), as it also drove high expression in the pharynx located very close to the XXX cells, and three did not show detectable expression in XXX cells (in adults; *mab-9, twk-39* and *lgc-36*). The absence of expression in the latter group could indicate that these genes represent false positives in the RAPID analysis, that the transcription was active only at earlier developmental timepoints, or that the arbitrarily defined promoters failed to reflect the endogenous transcriptional activity. Even if the subset of genes tested with the reporter follow-up analysis is small, we could calculate a gross estimate of the true positive rate of RAPID in XXX cells. Based on the 8/11 ratio (8 positive genes/promoters out of 11 interpretable readouts), a resampling bootstrap simulation indicated that the true positive rate should range between 45 and 88% (95% confidence interval, Figure S10). By extrapolation, this would correspond to a range of about 1100 to 2100 genes expressed in XXX and identified via RAPID.

**Figure 4.**
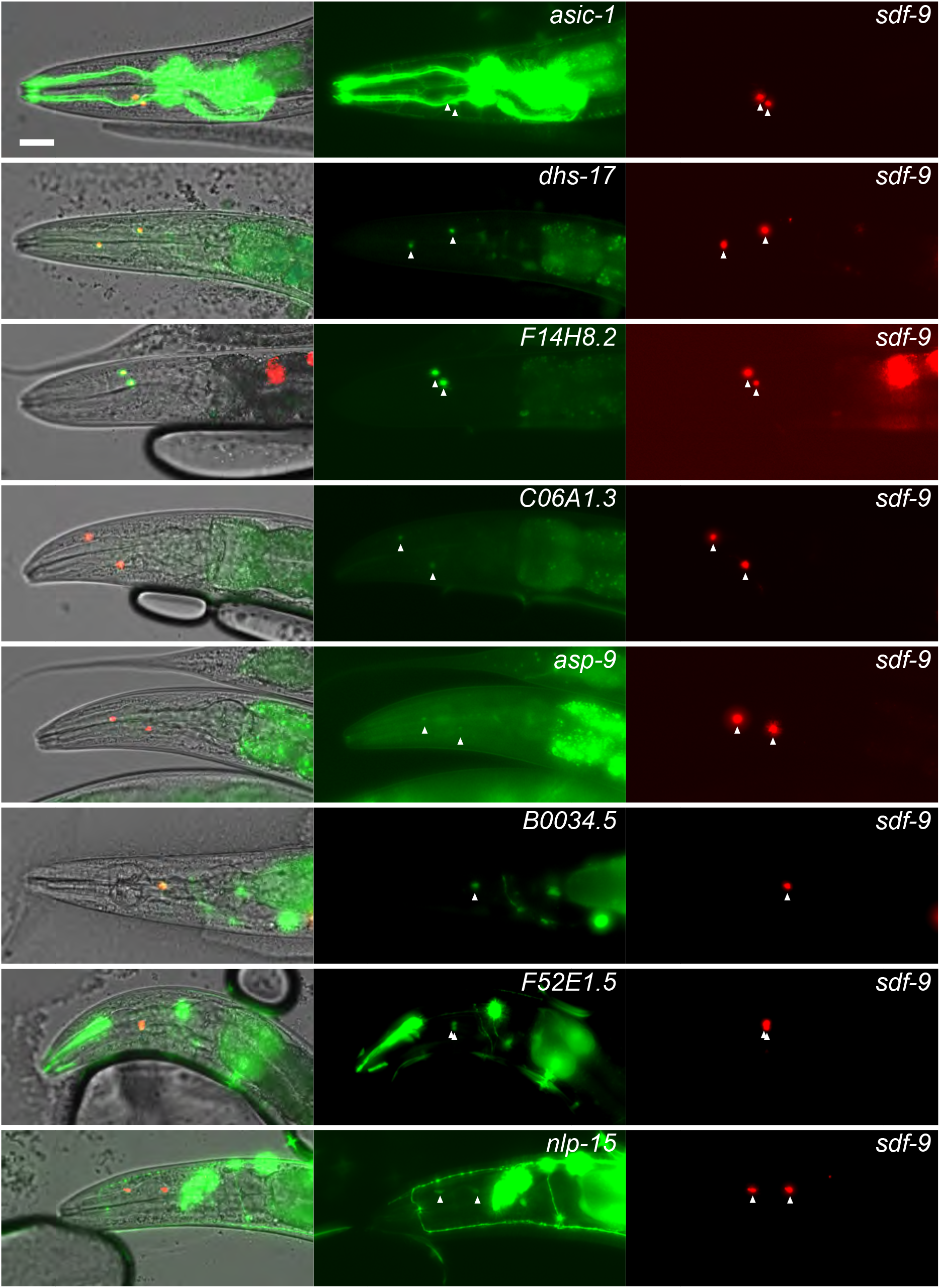
Promoters of genes identified as expressed in XXX cells using RAPID drive fluorescent reporter expression in XXX cells. Left panel, partial z-projections of adult heads showing expression of *sdf-9p::mScarlet* in XXX cells. Middle panel, partial z-projections of adult heads showing the expression pattern of *mNeonGreen* reporter under transcriptional control of the promoters of indicated genes. Right panel, merge of fluorescent channels within DIC images validating the expression of mentioned genes in XXX cells. Bar: 10 μm.

Interestingly, in half of the reporters expressed in XXX cells (4/8), we observed only limited or no signal in other cell types. In particular, reporters for *dhs-17* and *F14H8.2* were predominantly expressed in XXX cells and sequence homology suggests plausible XXX-specific roles for these two genes. Indeed, *dhs-17* encodes an uncharacterized member of the short-chain dehydrogenase/reductase family (SDR), which is potentially involved in steroid metabolism (Zhang *et al*. 2013) and may therefore contribute to this specific neuroendocrine function of XXX cells. As for *F14H8.2*, it is a paralog of *eak-4* selectively expressed in XXX cells and regulating dauer formation (Hu *et al*. 2006).

Taken together, the results of our reporter gene analysis indicate that the RAPID approach applied to only two cells per animal successfully identifies actively transcribed genes in these cells, including genes with relatively low expression and novel cell-specific markers.

## Discussion

Determining the transcriptional profile of specific cell types has been a major hurdle for *C. elegans* researchers, as most research is carried out on intact, living animals. Recent technological advances now allow the acquisition of a comprehensive view of the genes expressed in individual cell types. Two main RNA-based approaches have been used in the community to this aim:

- “RNA tagging/pulling” approaches that allow to sample gene expression directly from whole, live animals, either through affinity purification or pull-down enrichment.
- “dissociation-based” methods, requiring tissue dissociation followed by either purification of cells/nuclei by FACS or individual cell isolation.

Here, we adapt a third type of approach originally developed in *Drosophila*, based on RNA polymerase footprinting on DNA (Southall *et al*. 2013). RAPID is based on DNA modification by DNA adenine methyltransferase fused with a RNA polymerase subunit, detecting active transcribed genes footprinting by DamID (van Steensel *et al*. 2001). RAPID is a simple, fast and cost-effective technique to identify transcribed genes in individual tissues or cells, requiring only the knowledge of one tissue- or cell-specific promoter. We discuss below the specificities and limits of RAPID for transcriptional profiling, in comparison with previously published methods (summarised in Table S7).

First, as RAPID is based on the extraction of DNA from entire animals, it avoids *in vitro* cell manipulations and cell isolation-induced transcriptome modifications, similarly to RNA-tagging/pulling techniques. Second, since methylation occurs only in cells in which the Dam fusion is expressed after *Cre/lox* recombination, RAPID yields signals with a high specificity toward targeted tissues. Our comparative analyses with other methods for muscle and intestine, as well as the validation of our results with XXX cells, indeed indicate that a majority of RAPID hits are genuinely expressed in these tissues. Third, the very low level of signal background noise in RAPID makes it sensitive enough to function with a low number of worms (4’000, at least a 10-fold reduction compared to other methods), even for a tissue representing 2 cells per animal. This reduces the time necessary for worm population growth. Additionally, as each examined tissue only requires a tissue-specific Cre driver (many of which already exist; Kage-Nakadai *et al*. 2014; Ruijtenberg and van den Heuvel 2015), and as the library preparation is relatively simple to execute, RAPID makes it possible to process several conditions/tissues in parallel and is accessible to a neophyte researcher in the field. Fourth, RAPID is versatile, as minimal modifications will be required to target other DNA interacting proteins such as transcription factors or chromatin modifiers, in order to analyze tissue- or cell-specific genome-wide binding (Marshall and Brand 2017; Aughey *et al*. 2018). With the lower cost of nanopore sequencing systems, it is even conceivable that individual labs purchase their own equipment, further cutting down on waiting time and indirect costs.

The advantages described above come at a cost, in particular on how quantitative, dynamic and comprehensive RAPID data is. Our comparison of the overlap between transcriptomic datasets from different methods suggests that a certain portion of genes is uniquely detected by each method (Fig. S5), an indication that no single approach truly captures the complete transcriptome or that variable experimental conditions impact on the genes identified as expressed. However, when compared to all other RNA-seq based methods (Fig. S5), RAPID identified overall less genes as expressed. On the one hand, it is possible that affinity purification or dissociation-based methods could detect a number of genes due to contaminants and experiment-induced gene expression, or stored messengers inherited from mother cells. On the other hand, RAPID has a number of inherent technical limitations, which may hinder a comprehensive and unbiased genome interrogation, as described below.

1. RAPID relies on the GATC density per gene, and a lower density will yield a lower signal, as longer *Dpn* I restriction fragments will be less efficiently amplified by the DamID PCR. In agreement, genes detected using FACS/RNA-seq but not by RAPID, have on average a lower GATC density (data not shown and Kaletsky *et al*. 2018). Increasing the material amount and/or sequencing coverage is expected to dampen this type of bias.
2. Another consequence of the methylation is that the dynamic range of RAPID compared to RNA-seq is expected to be lower: once a gene sequence is fully methylated, further transcription will not lead to increased methylation levels of the DNA, leading to a plateau effect. Conversely, as RAPID uses minute traces of the Dam fusions, lowly expressed genes will rarely be bound by a polymerase containing the Dam fusion, and their sequence will be rarely methylated. As our quantitative comparison reveals (Fig. 1C/3B), these issues are partially relieved by the stochasticity of the methylation between different cells in different animals, leading to a correlation between RAPID levels and expression levels determined by RNA-seq. Increasing sequencing coverage should allow both the retrieval of more lowly expressed genes and an improvement in the signal dynamic range. For certain applications, such as identifying tissue-specific genes to serve as molecular markers, a lower dynamic range can actually be an advantage as compared to RNA-seq methods, as the large range of mRNA molecule number in a cell requires very deep sequencing in order to detect more lowly expressed genes.
3. RAPID signal on DNA will not discriminate between gene isoforms created by alternative splicing, since the RNA polymerase will progress over the whole intronic and exonic regions regardless of whether they are retained in the final mRNA product.
4. RAPID signal is a stochastic average of the methylation by polymerases since the last DNA replication (which erases the methylation signal). The comparison between embryonic blastomeres and young adult tissues (with older post-mitotic cells) highlights the increase in reproducibility of RAPID as animals age (Fig. S4). Genes detected by RAPID are likely to represent cell-specific genes which are expressed over longer time scales in the cells under study, rather than transient expression in response to the environment. While this phenomenon enables RAPID to efficiently identify cell type-specific genes (Tables S4,S5), it will also limit its ability to report quantitative variations over shorter timescales.
5. In its present form, RAPID is not well suited to analyse transcriptome dynamics, as the methylation signal can only be erased by DNA replication. RAPID could be further improved by timing the expression of the Dam fusions. We tested auxin-mediated Dam degradation. However, degradation was not complete enough and the remaining low levels of Dam fusions were sufficient to create a RAPID signal (data not shown). Alternatively, degron-tagged Cre recombinases might provide a more reliable way to time the expression of the Dam. In a post-mitotic context however, RAPID should be able to identify transcriptional up-regulation events in response to environmental changes, as successfully achieved in *Drosophila* (Widmer *et al*. 2018). In addition, recent publications have highlighted the existence of a large cohort of genes with oscillating expression between molts (Kim *et al*. 2013; Hendriks *et al*. 2014; Meeuse *et al*. 2020; Hutchison *et al*. 2020). RAPID is unlikely to capture this type of phenomenon, which could turn into a desirable feature when one wants to mitigate this variability, *e.g*. in studies comparing hard-to-synchronize genetic backgrounds.

In conclusion, we believe that RAPID is a useful addition to the existing methods to analyse comprehensive gene expression at the cell type level, allowing one to identify new and specific markers and further study their biology. RAPID is an easy, scalable, entry-level method to target rare or less known cell types, cell types difficult to purify because of their morphology or the developmental stage targeted or when synchronisation of large populations is tedious, notably when multiple genotypes have to be assessed.

## Acknowledgments

We are grateful to Laurence Bulliard, Cihan Enci and Lisa Schild for expert technical support; Marc Hammarlund, Piali Sengupta and Sanders van der Heuvel for the gift of material; Michalis Barkoulas for sharing preliminary data; some strains were provided by the CGC, which is funded by NIH Office of Research Infrastructure Programs (P40 OD010440). The study was supported by the Swiss National Science Foundation (IZCNZ0_174703/SBFI_C16.0013, BSSGI0_155764 and PP00P3_150681 to D.A.G.; ZCSZ0-174641/SBFI_C15.0076, PP00P3_159320 and 31003A_176226 to P.M.), the ERC CoG PlastiCell grant #648960 to S.J. and a BMBS COST Action (BM1408).

## Supplementary figures

**Figure S1.**
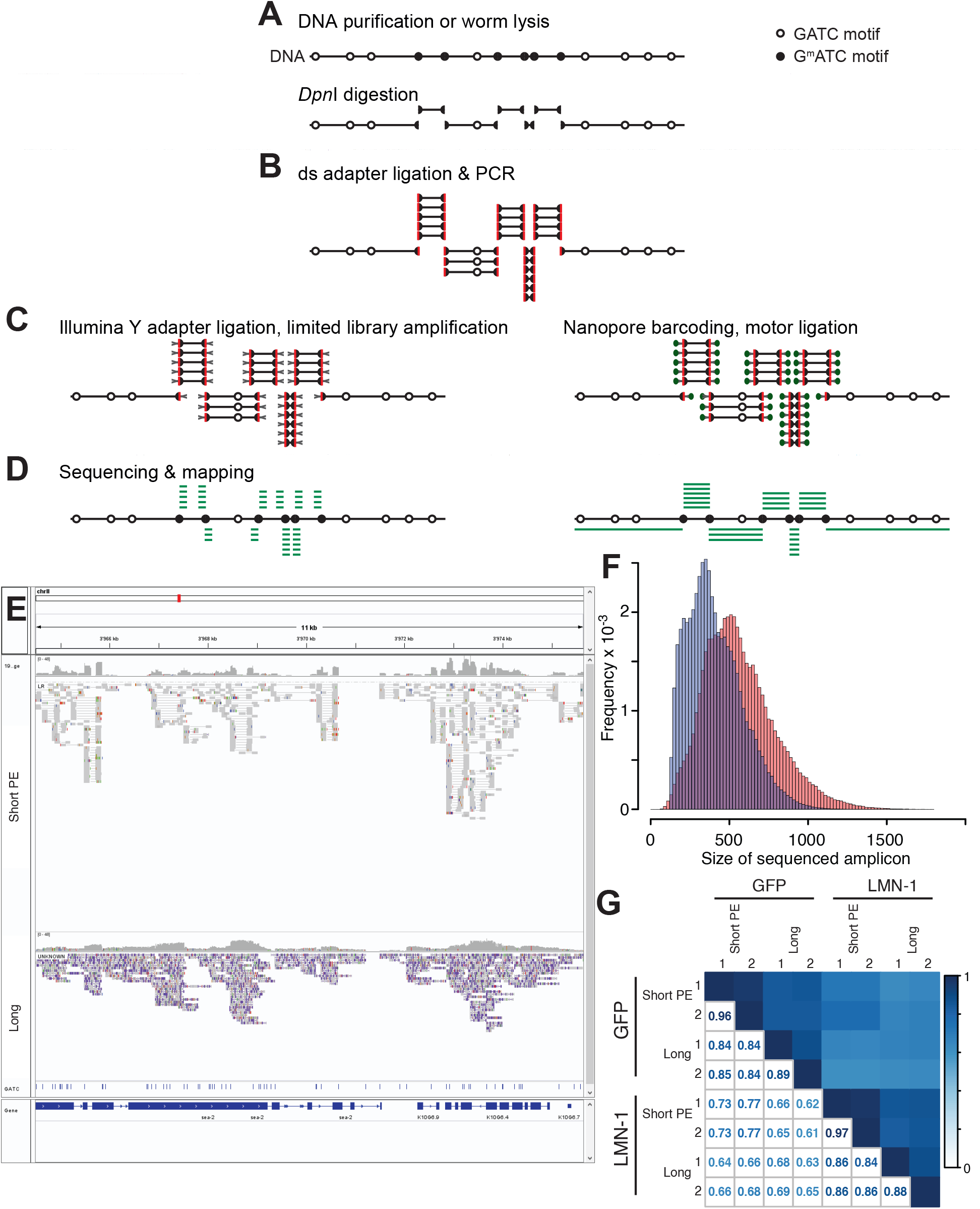
DamID Sequencing using Illumina or Nanopore technologies. **A.** Principle of methylated GATC identification using DamID. Methylated GATC motifs are cut using the methylation-sensitive *Dpn*I restriction enzyme, before **B.** adapter ligation and PCR amplification using a primer hybridizing in the adapter. **C.** For Illumina sequencing, classical DNA library preparation is used, involving Y adapter ligation and limited PCR amplification. DamID amplicons can only be sequenced on non-patterned Illumina flow cells as their variable length is detrimental to cluster formation and polony amplification. To solve the latter issue, amplicons from the first PCR round were directly sequenced using ligation-mediated library preparation and ONT long read sequencing. **D.** While Illumina sequencing highlights the beginning of the amplicons (and the end in case of paired end sequencing), ONT long read sequencing captures the entire length of the amplicon. **E.** Snapshot of the same library sequenced with Illumina short read sequencing (paired end, upper panel) and ONT long read sequencing (bottom panel). **F.** Comparison of amplicon length obtained from both sequencing techniques for the same library (blue, Illumina; orange, ONT). The smaller sizes of Illumina-sequenced amplicons is due to the loss of these amplicons during polony amplification in Illumina sequencing, as the average size of the amplicons determined using automated electrophoresis before loading onto the flow cell correspond to the ONT-sequenced one (data now shown). **G.** Pearson correlation at the single GATC fragment resolution for 4 different libraries of 2 different Dam fusions (with either GFP or LMN-1) sequenced with either Illumina (short PE) or ONT (Long) sequencers.

**Figure S2.**
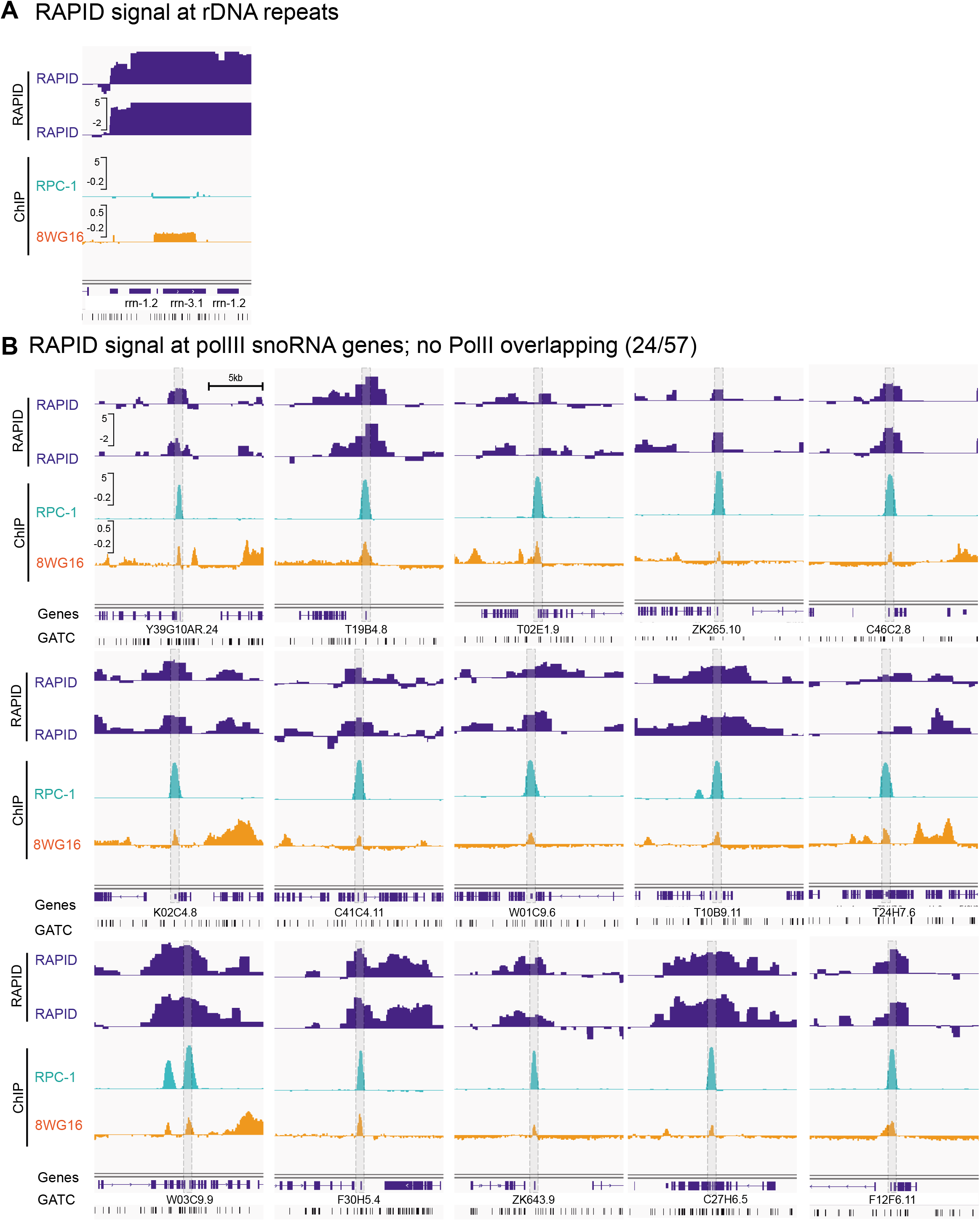
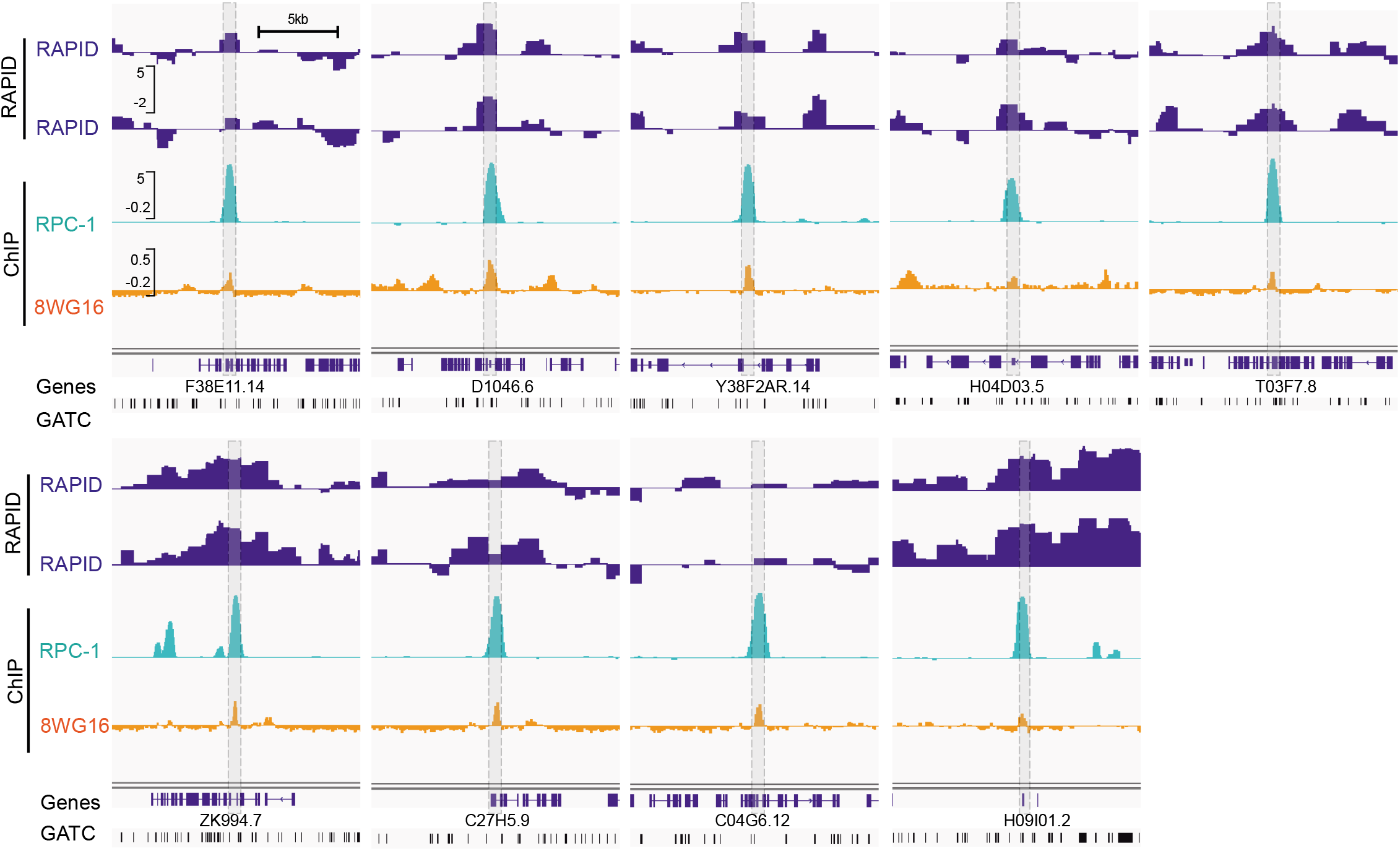
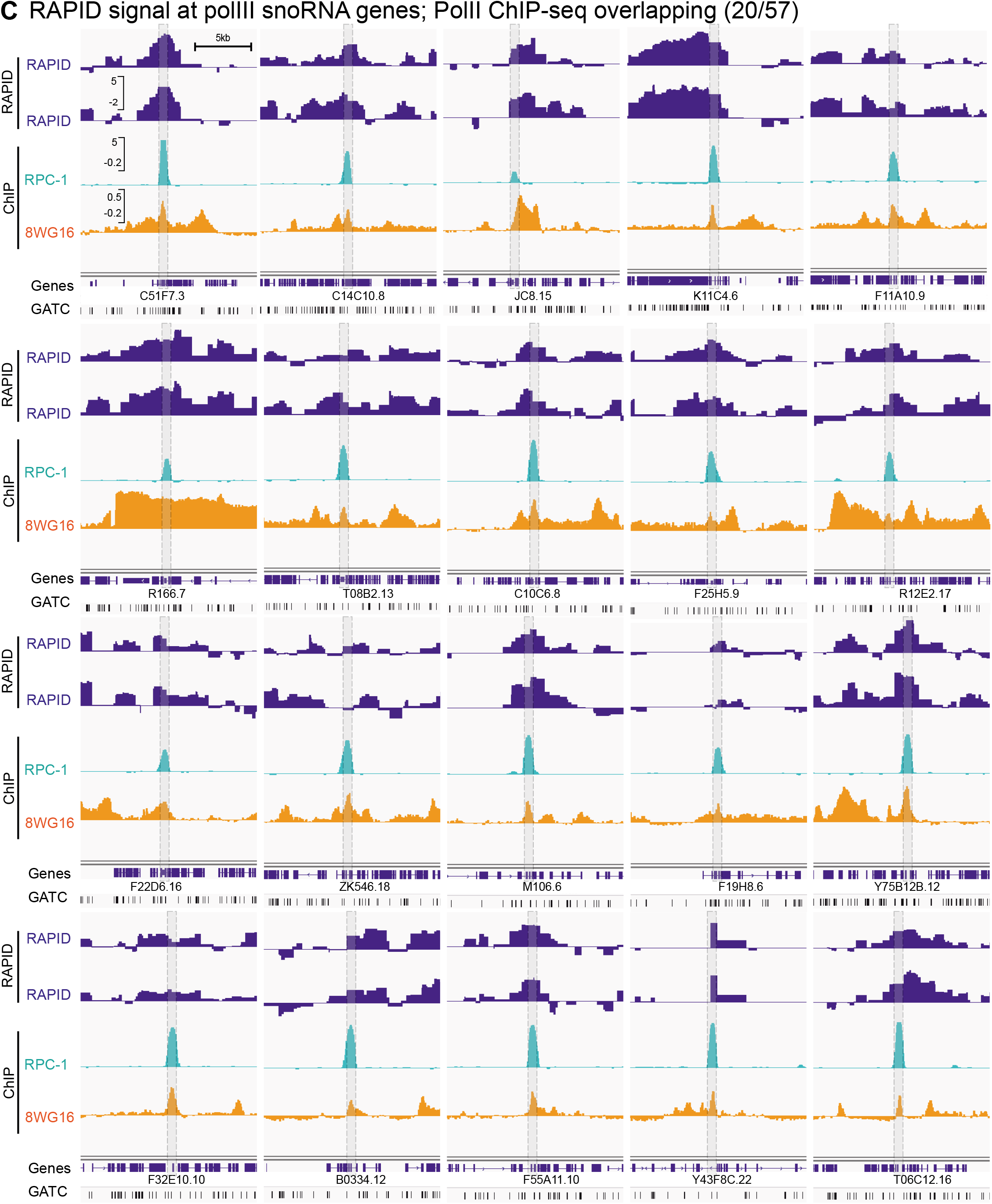
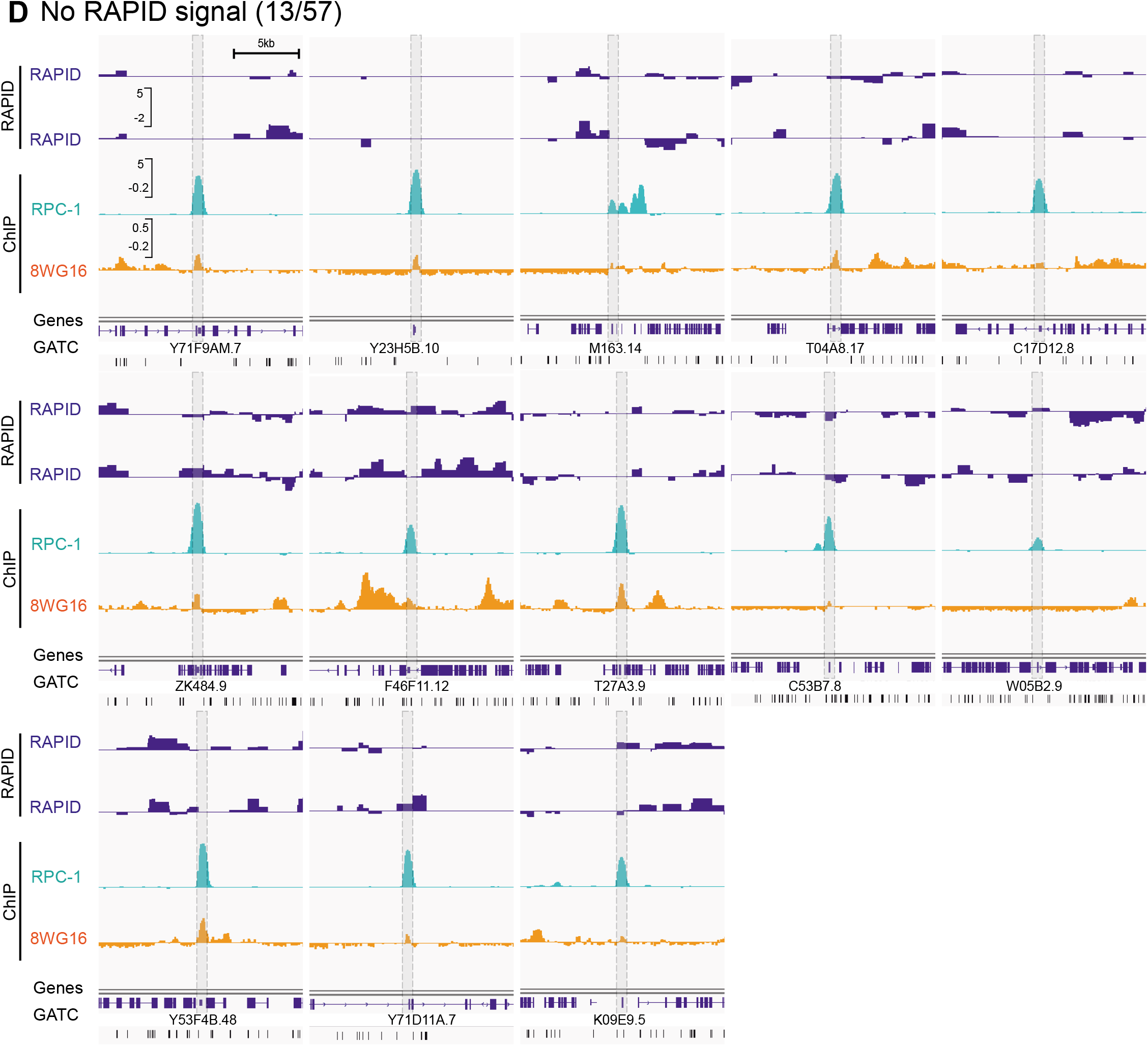
RAPID allows identification of PolI and PolIII transcribed genes. **A.** RAPID footprinting of the rDNA locus on the right telomere of chromosome I, together with RNA polymerase II ChIP-seq (8WG16 ChIP-seq from young adult animals from (Miki *et al. 2017*). **B.** RAPID signal at PolIII snoRNA genes, with no PolII overlapping signal (24 out of 57 genes; 8WG16 ChIP-seq as in A). PolIII-specific genes were taken from (Ikegami and Lieb 2013), ChIP-seq for RPC-1 (RNA polymerase III subunit A) performed in embryos. The 8WG16 antibody produces a low residual signal on the snoRNA genes. **C.** RAPID signal at polIII snoRNA genes with overlapping PolII ChIP-seq signal (20 out of 57 genes; data as in A and B). **D.** snoRNA genes with no RAPID signal (13 out of 57 genes; data as in A and B). Classifications were done twice independently by visual inspection of the tracks.

**Figure S3.**
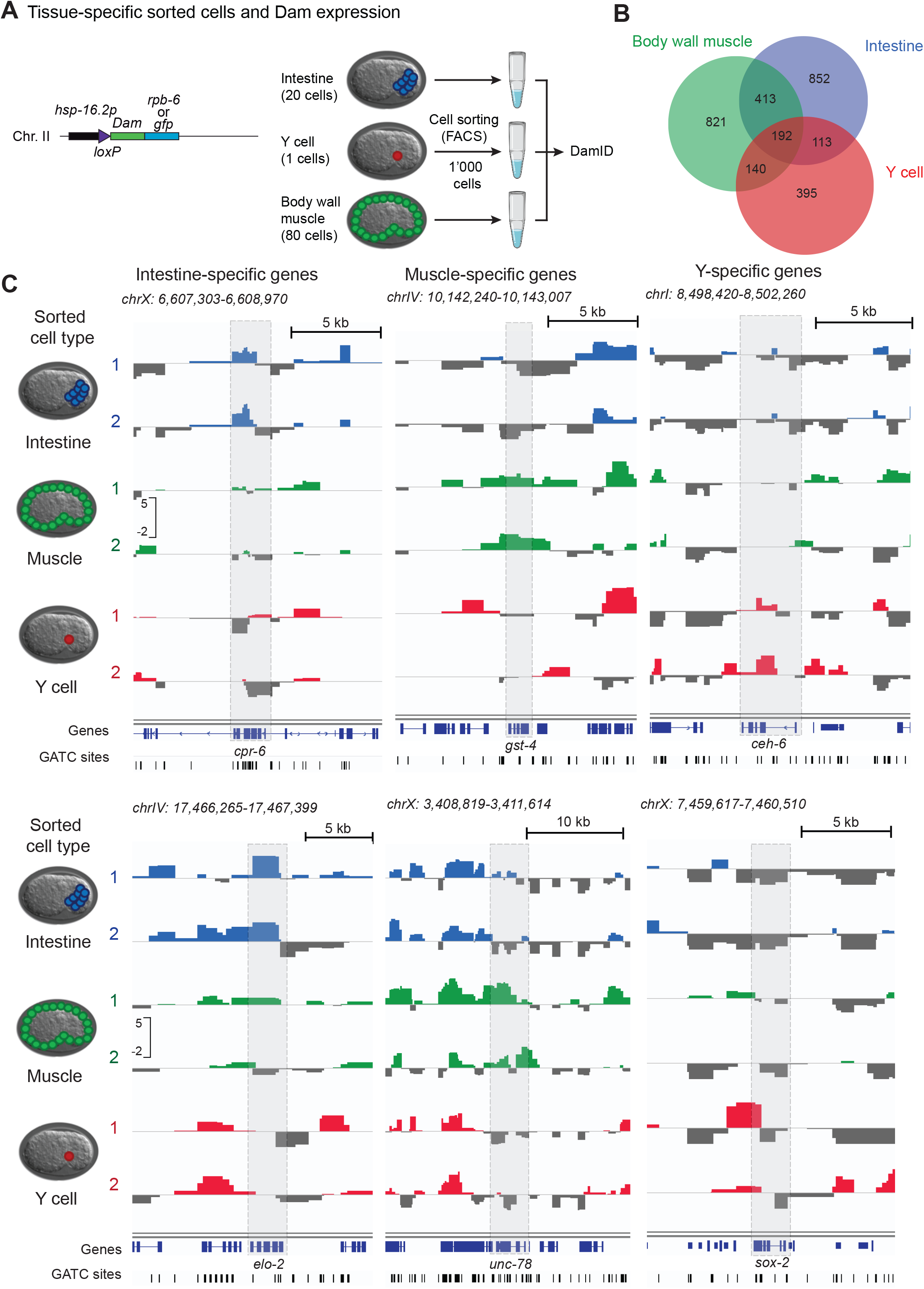
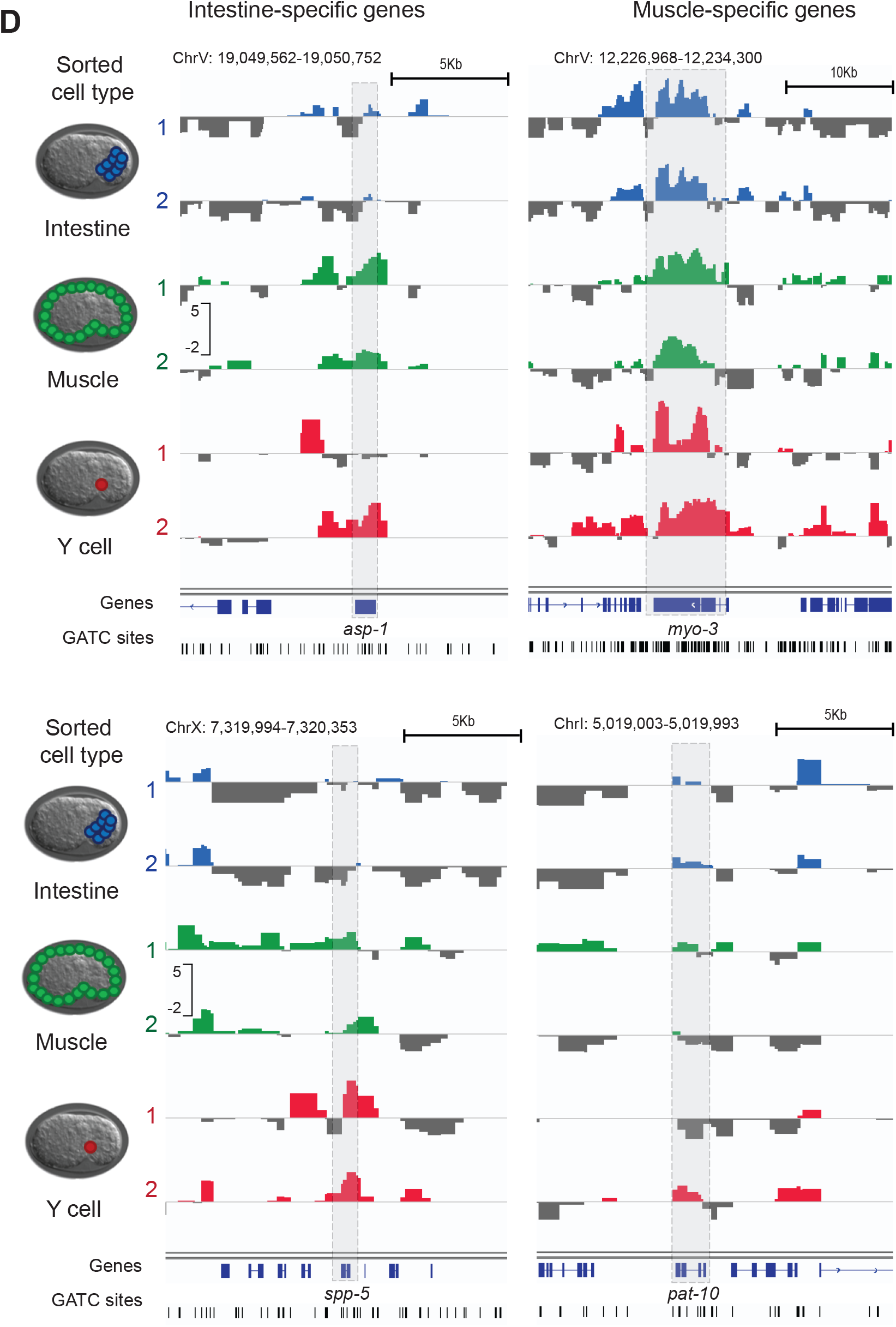
RAPID using sorted embryonic blastomeres. **A.** Procedure schematic. Embryonic blastomeres from a strain ubiquitously expressing trace amounts of Dam constructs (either fused to GFP or RPB-6) were sorted based on the expression of fluorescent markers using fluorescence-activated cell sorting (FACS). For each construct and each tissue, 1’000 cells were pooled before performing DamID. **B.** Venn diagram showing overlap between RAPID footprinted genes (FDR < 0.05 in both replicates). **C, D.** Individual profiles for genes specific for the different tissues tested (intestine, muscle and Y cell).

**Figure S4.**
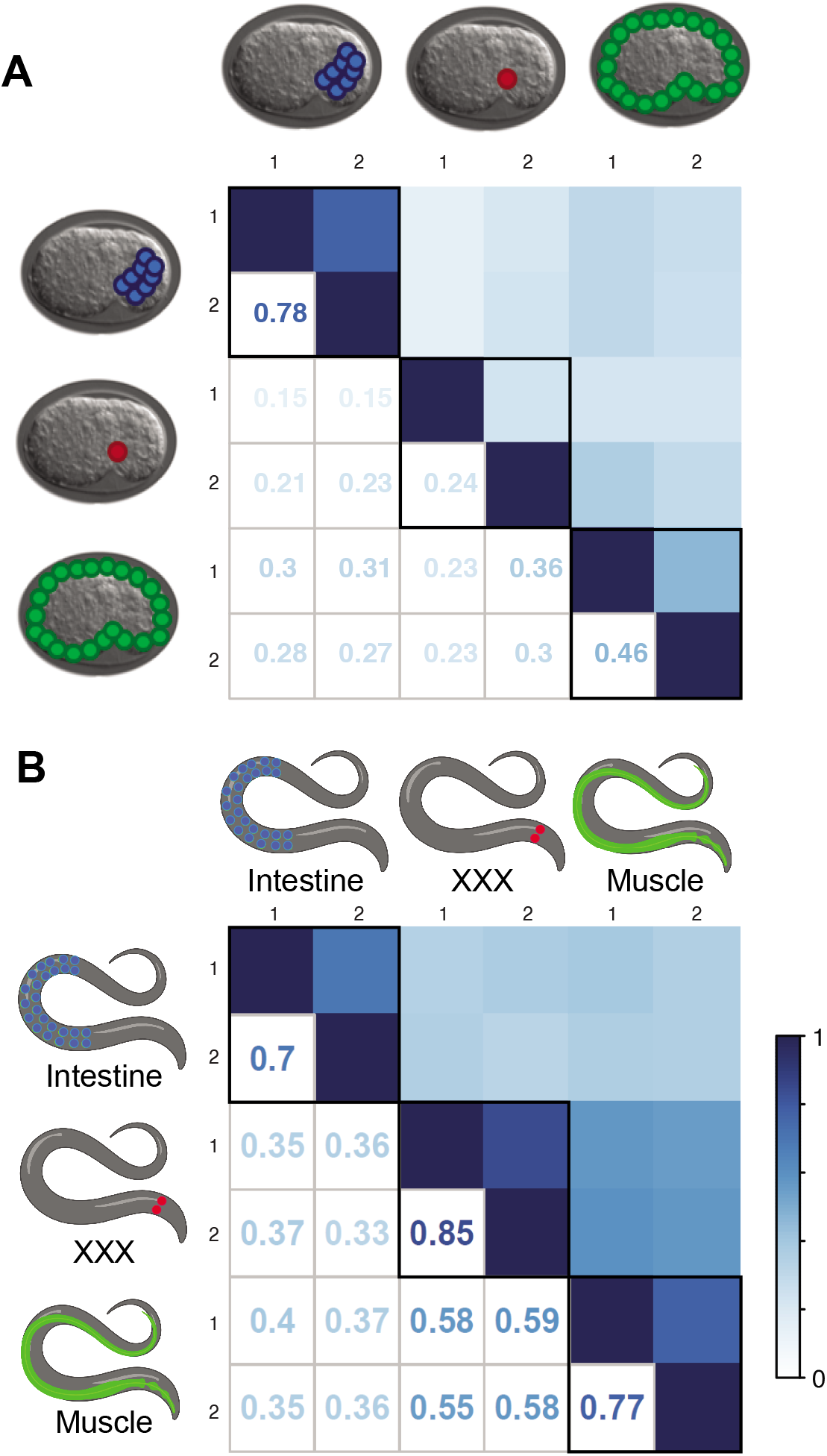
Pearson correlation coefficient between RAPID libraries at the single gene level resolution for the different tissues tested. **A.** Correlation of RAPID footprinting signal between replicates performed in different tissues in embryonic blastomeres sorted using fluorescent markers. **B.** Correlation of RAPID footprinting signal between replicates performed in different adult tissues using Cre-mediated recombination for tissue-specific expression of the Dam fusions.

**Figure S5.**
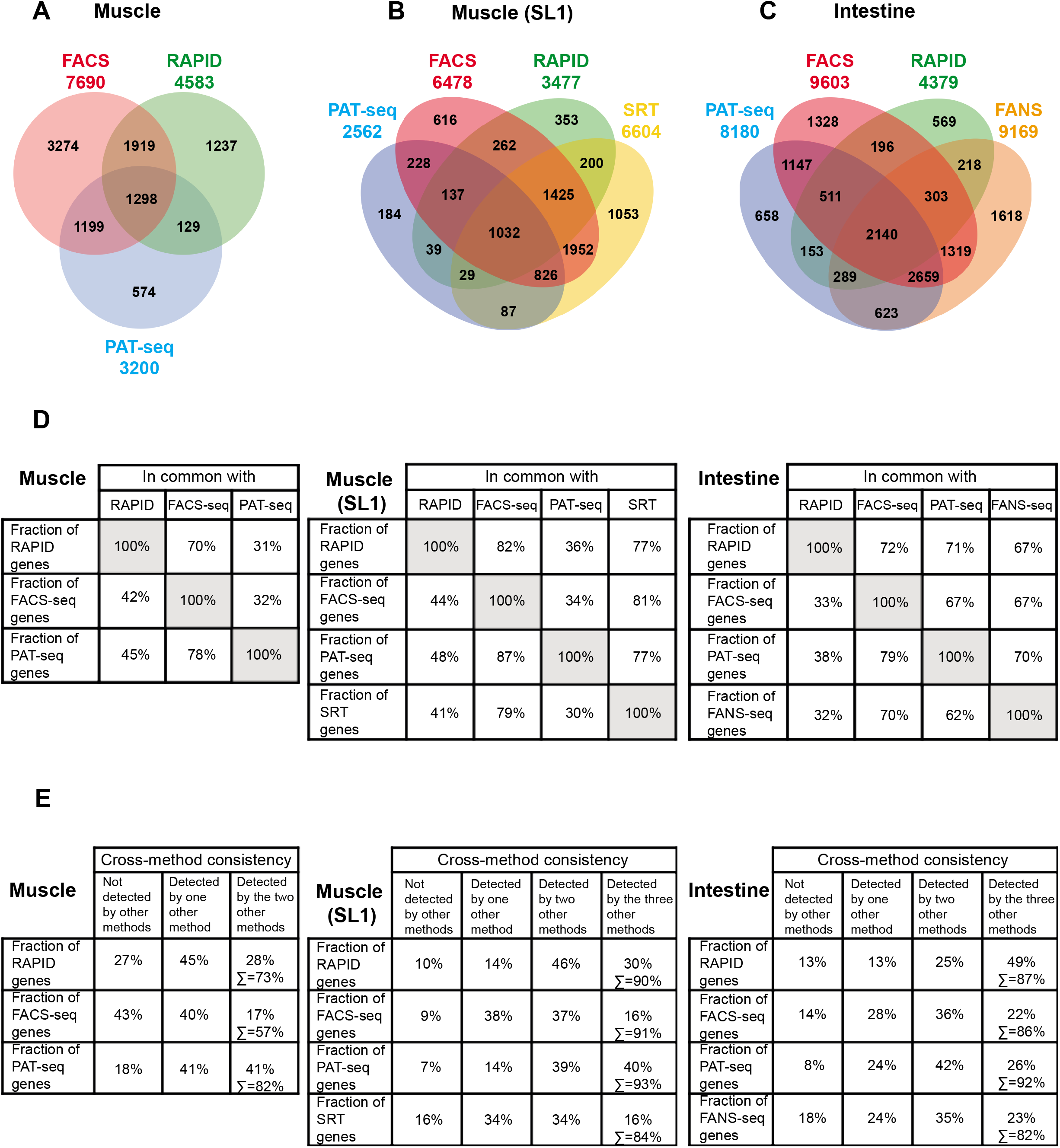
Extended comparative analysis of the muscular and intestinal transcript profiles obtained with RAPID and RNA-seq-based methods. **A.** Overlap of genes footprinted using RAPID with muscle-expressed genes detected by FACS-seq (Kaletsky *et al*. 2018) and PAT-seq (Blazie *et al*. 2017). **B.** Overlap of SL1-trans-spliced genes detected by different methods in muscle: RAPID (this study), PAT-seq (Blazie *et al*. 2017), FACS ((Kaletsky *et al*. 2018) and SRT (Ma *et al*. 2016). The selection of SL1-trans-spliced protein-coding genes was made according to the annotation of modENCODE (Allen *et al*. 2011, Ma *et al*. 2016). **C.** Overlap of genes footprinted using RAPID with intestine-expressed genes detected by PAT-seq (Blazie *et al*. 2017), FACS (Kaletsky *et al*. 2018) and FANS (Haenni *et al*. 2012). **D.** Tables showing the percentage of genes commonly detected by method pairs (listed in A-C, respectively). For the muscle transcriptome, two tables are shown: one with all the genes detected by the methods in A (left) and another with the SL1-trans-spliced genes as used for the analysis in B (middle). **E.** Cross-method consistency as assessed with the fractions of genes detected by one, two or three methods. The percentage of genes detected by at least one method is indicated in the last column (Σ). The analysis corresponding to muscle and muscle-SL1 is shown in the left and middle tables. The analysis corresponding to the intestine is shown in the table to the right.

**Figure S6.**
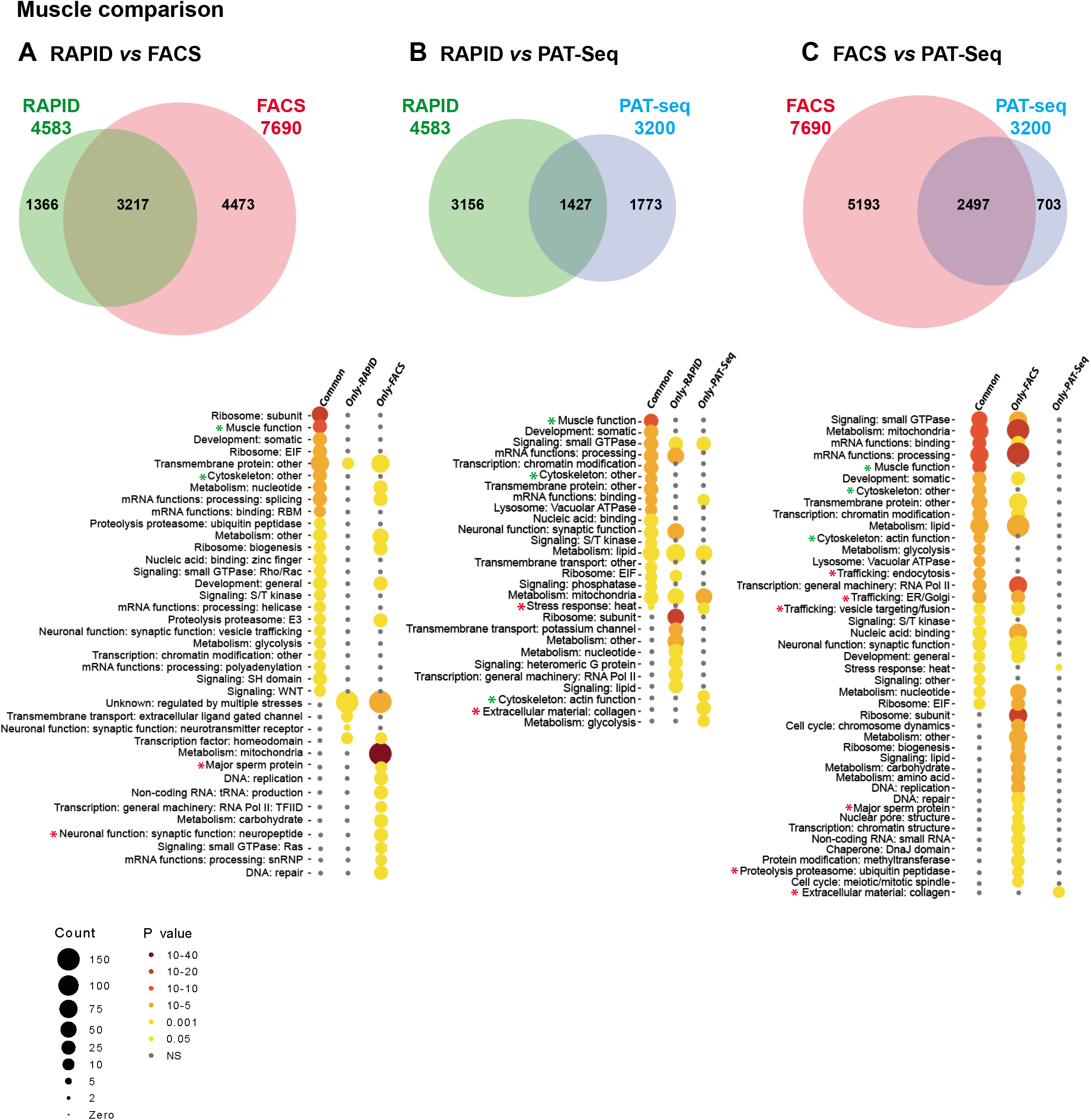
Muscle transcriptome GO-analysis. Comparison of gene sets commonly identified by two methods (Common) and gene sets identified by only one method (Only-RAPID, Only-FACS, Only-PAT-seq). **A.** Comparison between RAPID (this study) and FACS (Kaletsky *et al*. 2018). **B.** Comparison between RAPID (this study) and PAT-seq (Blazie *et al*. 2017). **C**. Comparison between FACS ((Kaletsky *et al*. 2018) and PAT-seq (Blazie *et al*. 2017). Functional enrichment analysis was performed using WormCat. The pervasively represented category “unknown” was removed from the depicted heatmap. Green asterisks highlight expected muscle-specific categories and red asterisks highlight less expected categories for this tissue (more typical in other tissues; Holdorf *et al*. 2020).

**Figure S7.**
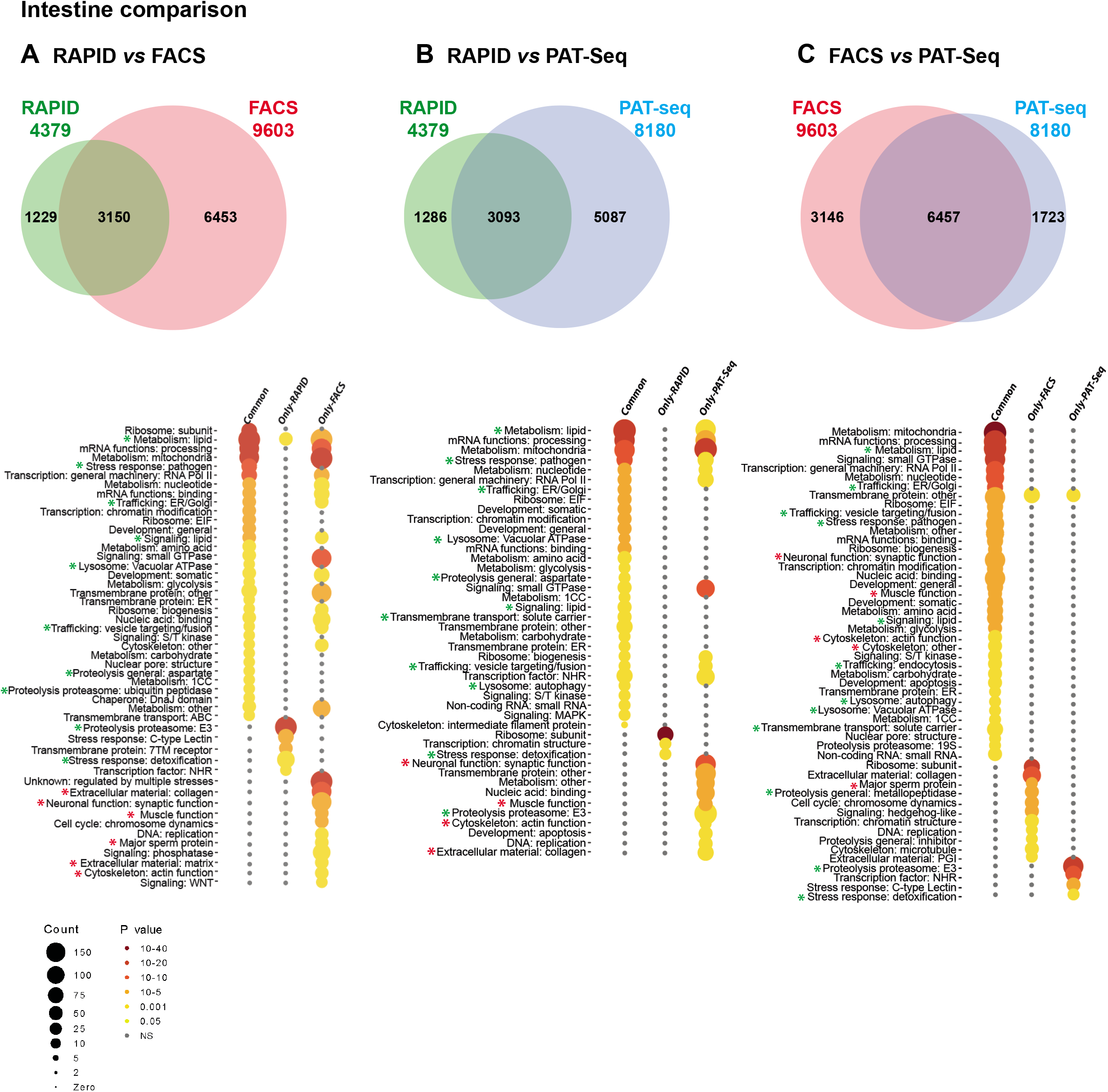
Intestine transcriptome GO-analysis. Comparison of gene sets commonly identified by two methods (Common) and gene sets identified by only one method (Only-RAPID, Only FACS, Only PAT-seq). **A.** Comparison between RAPID (this study) and FACS (Kaletsky *et al*. 2018). **B.** Comparison between RAPID (this study) and PAT-seq (Blazie *et al*. 2017). **C.** Comparison between FACS (Kaletsky *et al*. 2018) and PAT-seq (Blazie *et al*. 2017). Functional enrichment analysis was performed using WormCat. The pervasively represented category “unknown” was removed from the depicted heatmap. Green asterisks highlight expected intestine-specific categories and red asterisks highlight less expected categories for this tissue (more typical in other tissues; Holdorf *et al*. 2020)

**Figure S8.**
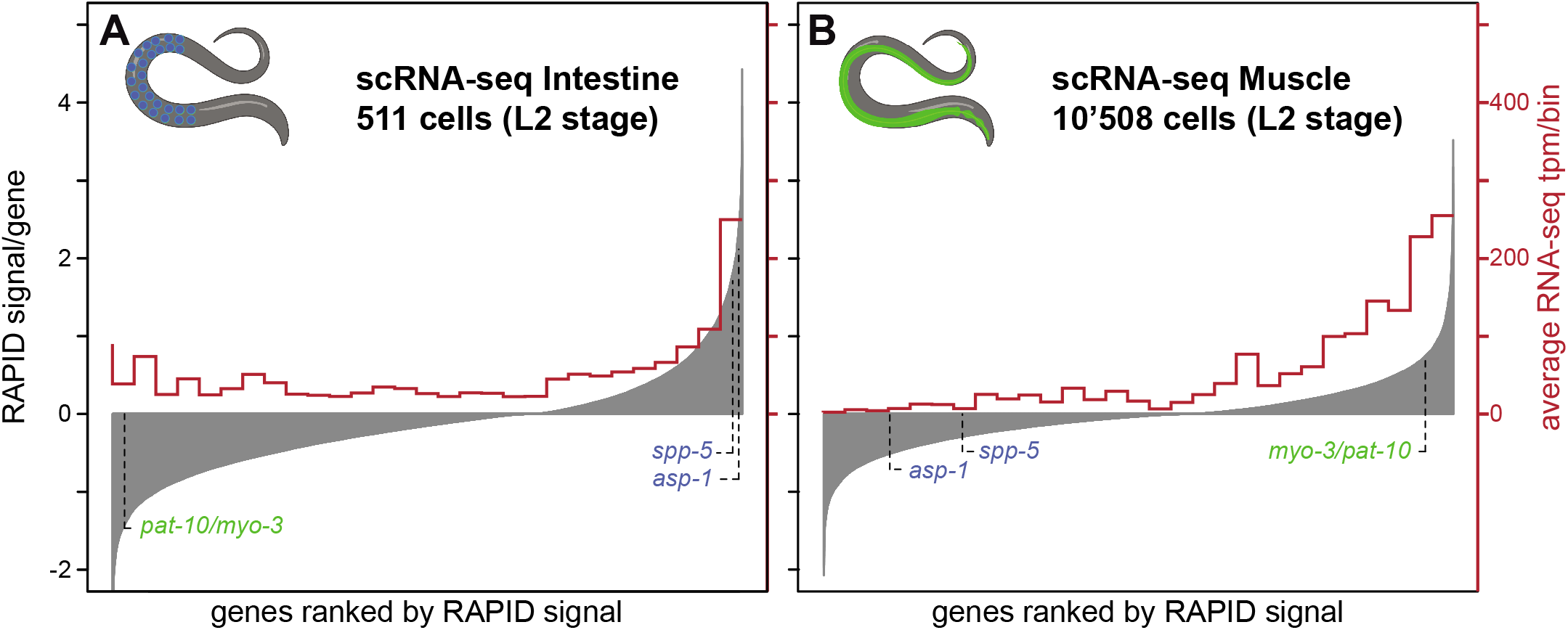
Semi-quantitative comparison between RAPID and sciRNA-seq (L2 larval stage; Cao *et al*. 2017) in intestine (**A**) and muscle (**B**). All genes are represented on the x axis ranked, from left to right, based on the RAPID signal in the considered tissue (shown on left y axis). Averages of transcripts per million (tpm) for those genes in cells identified as intestine (511 cells) or muscle (10’508 cells) were calculated, using the genes falling into each bin of 690 genes on the x axis (values on right y axis). The muscular and intestinal marker genes presented in Fig. 2C are indicated in green and blue, respectively.

**Figure S9.**
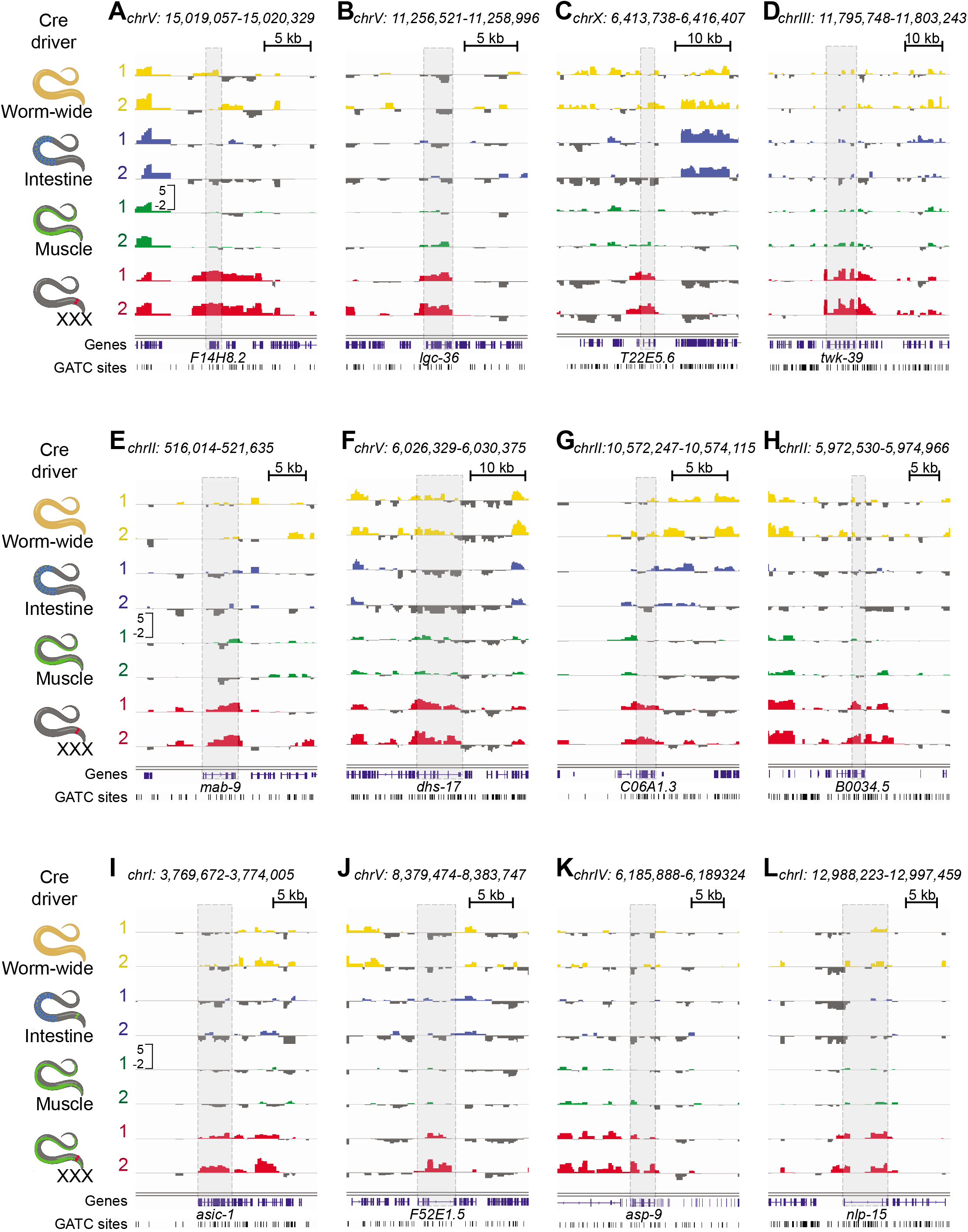
RAPID profile of candidate genes with XXX-specific expression. RAPID footprinting of twelve candidate genes expressed in a tissue-specific manner in XXX cells but not in muscle, intestine and enriched worm-wide are shown. Candidate XXX genes are ordered according to the RAPID footprinting level.

**Figure S10.**
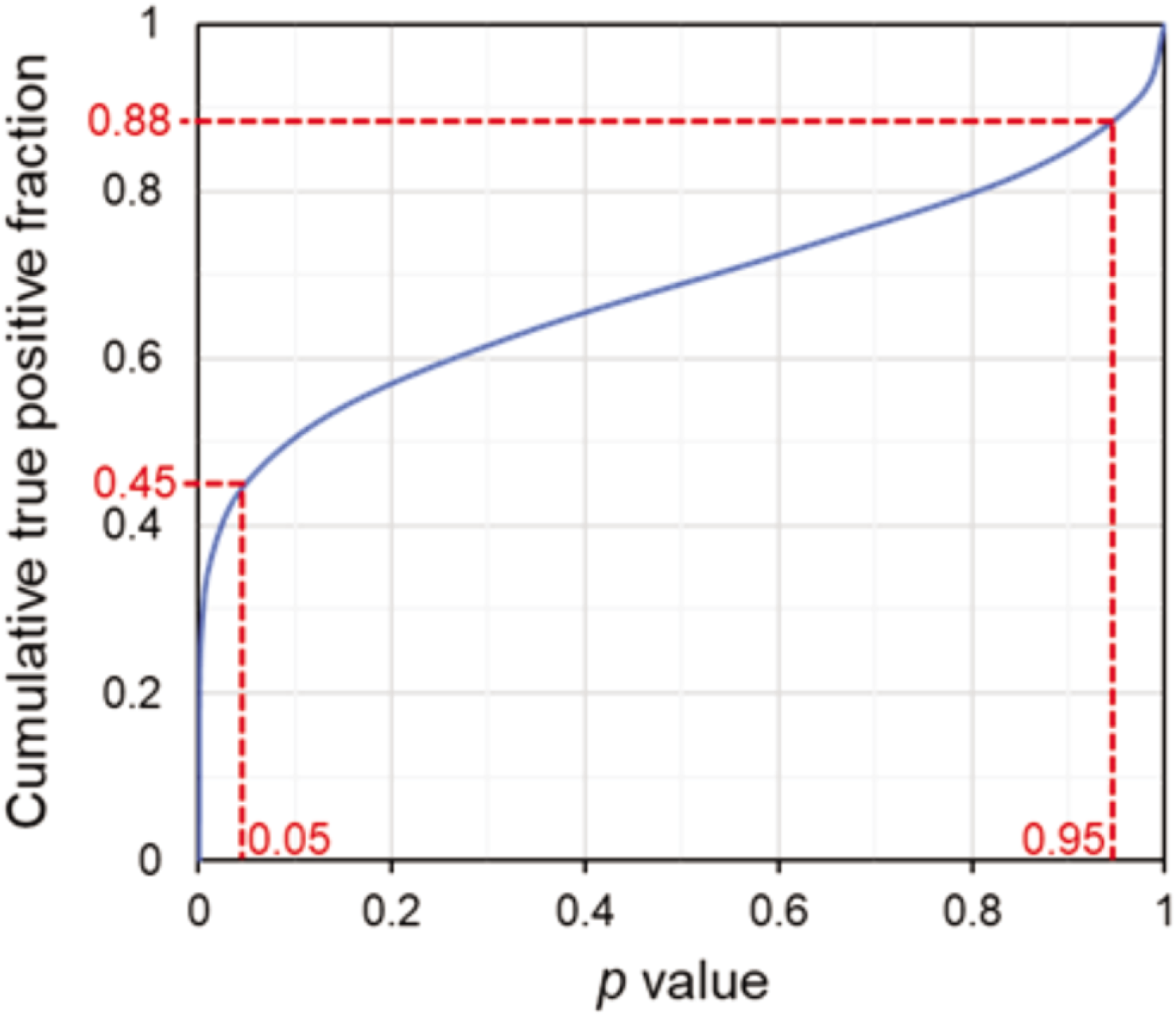
Estimation of the true positive rate in XXX-specific RAPID hits. Cumulative distribution of the predicted number of true positive hits in the XXX RAPID dataset, based on the fraction of true positives in the gene subset tested with the reporter gene approach. Resampling was performed with 10’000 bootstrap iterations. Boundaries of the 95% confidence interval are depicted in red.

## Supplementary information

**Table S1.**
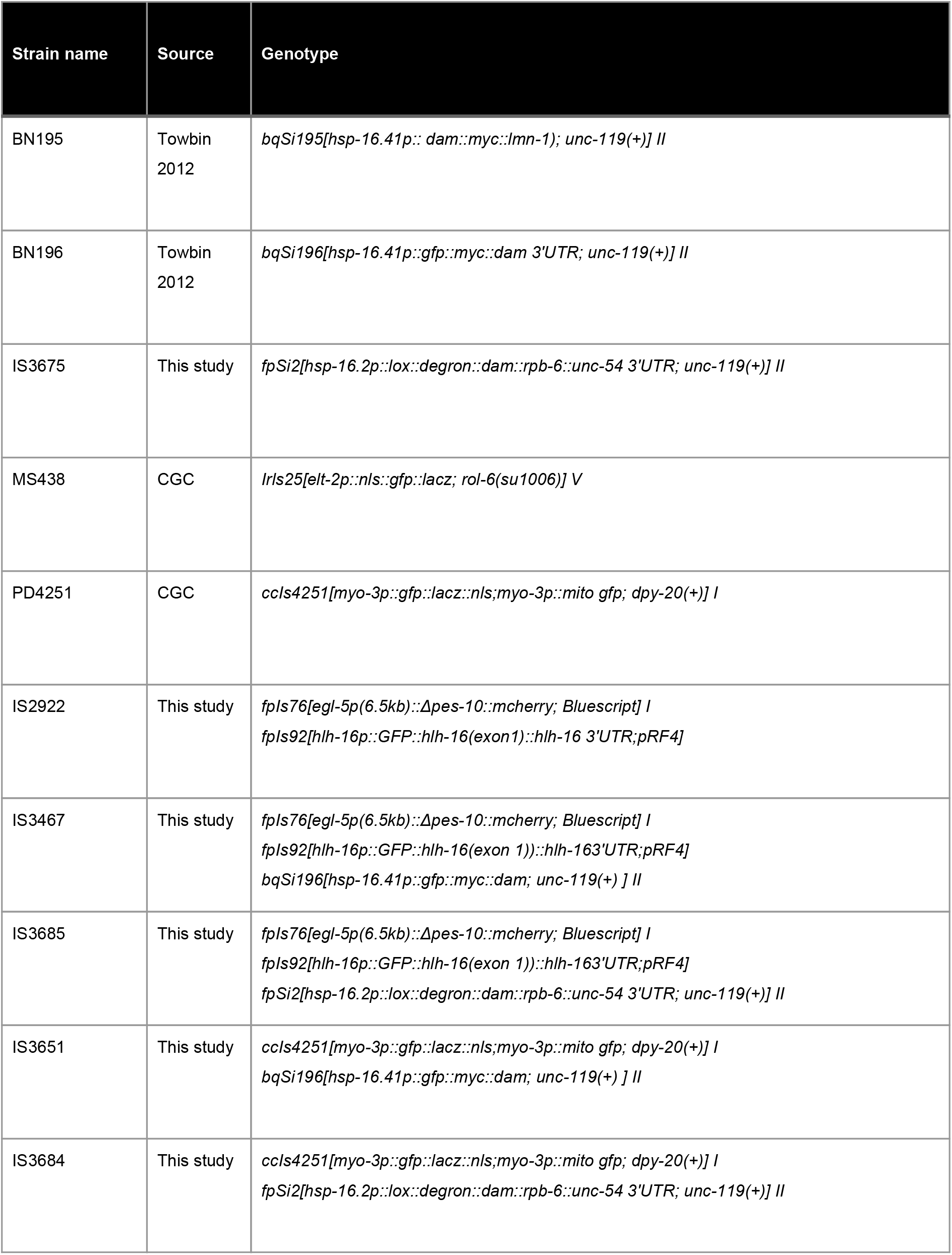
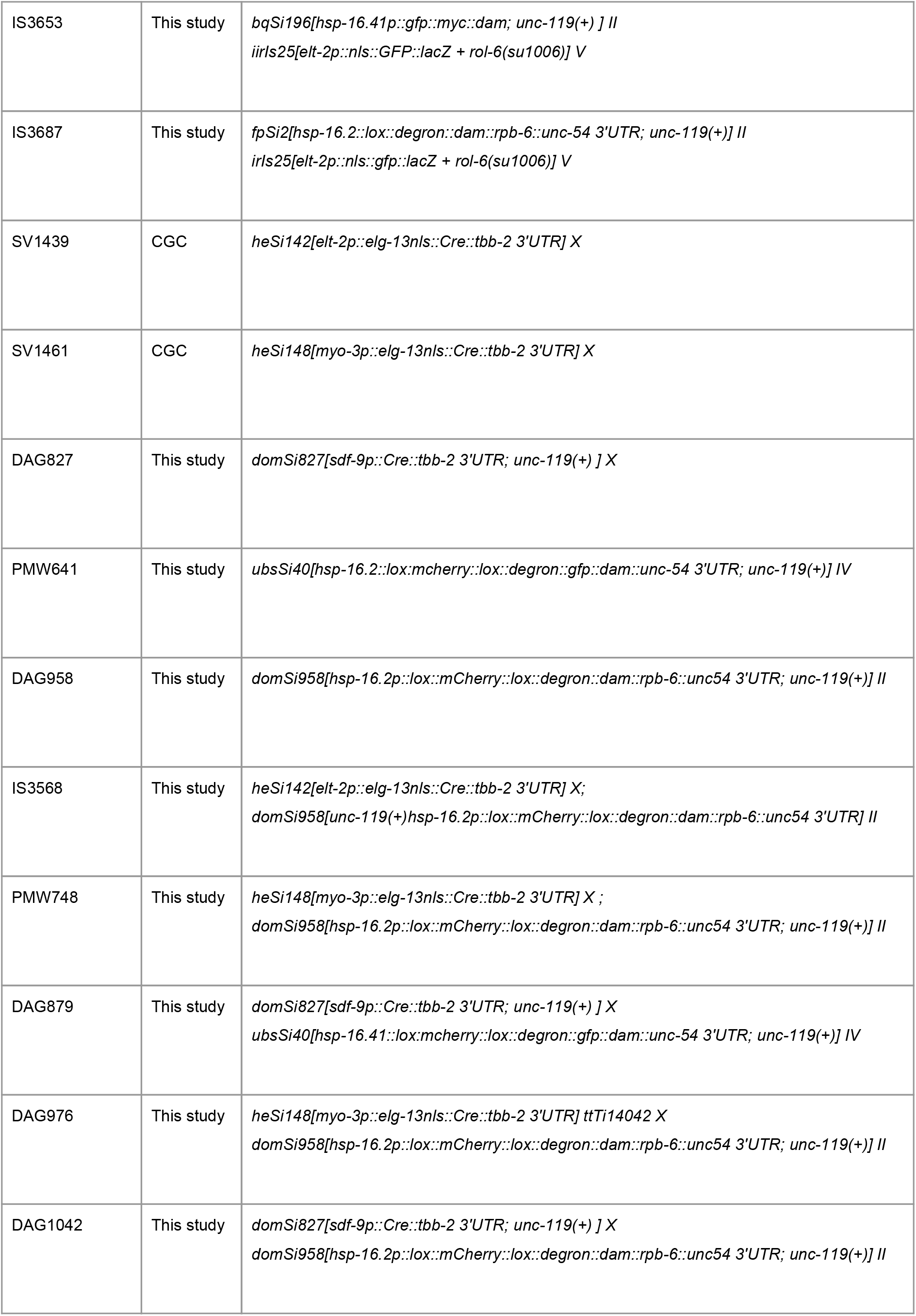
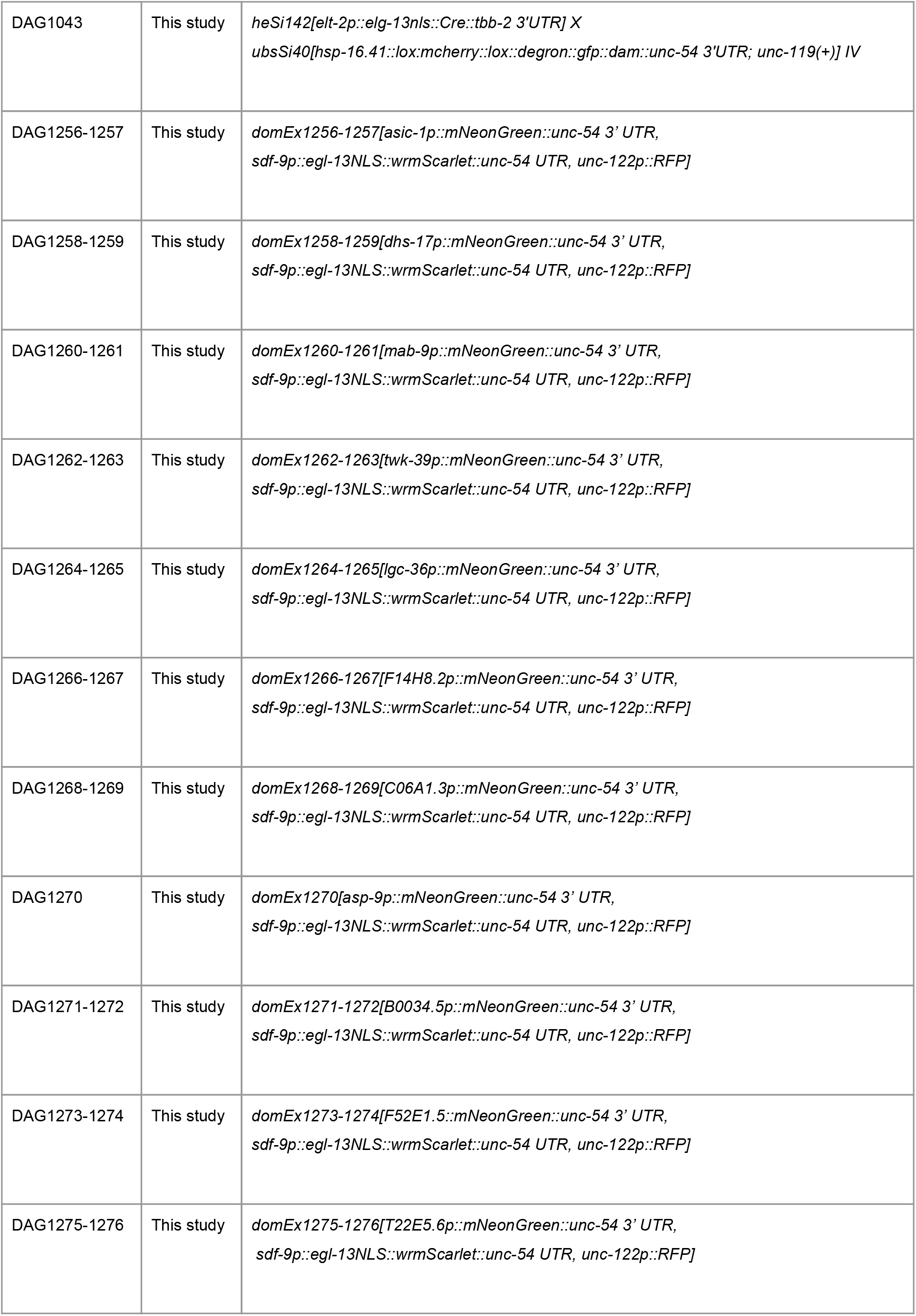
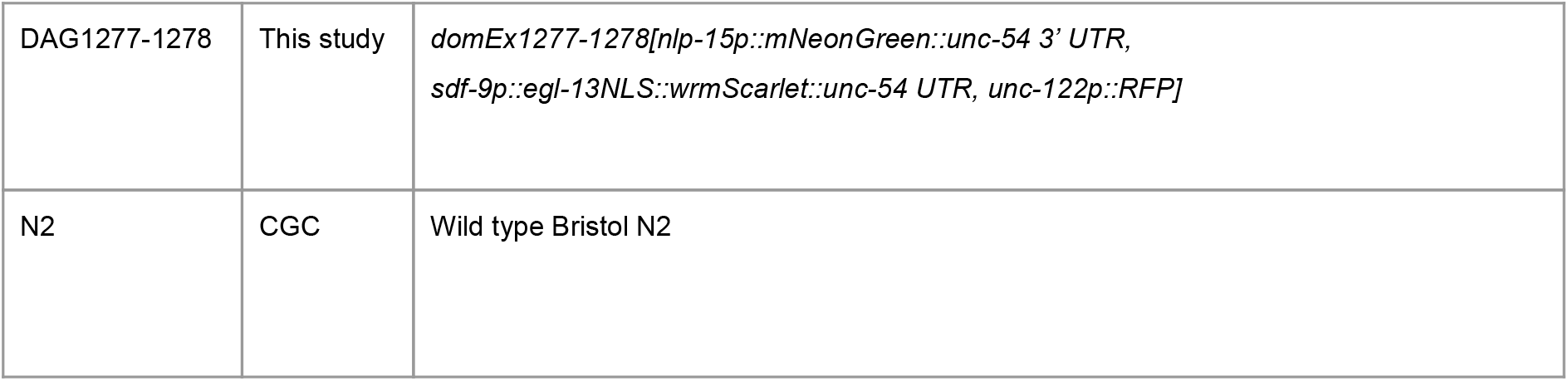
Worm strains collection

**Table S2.**
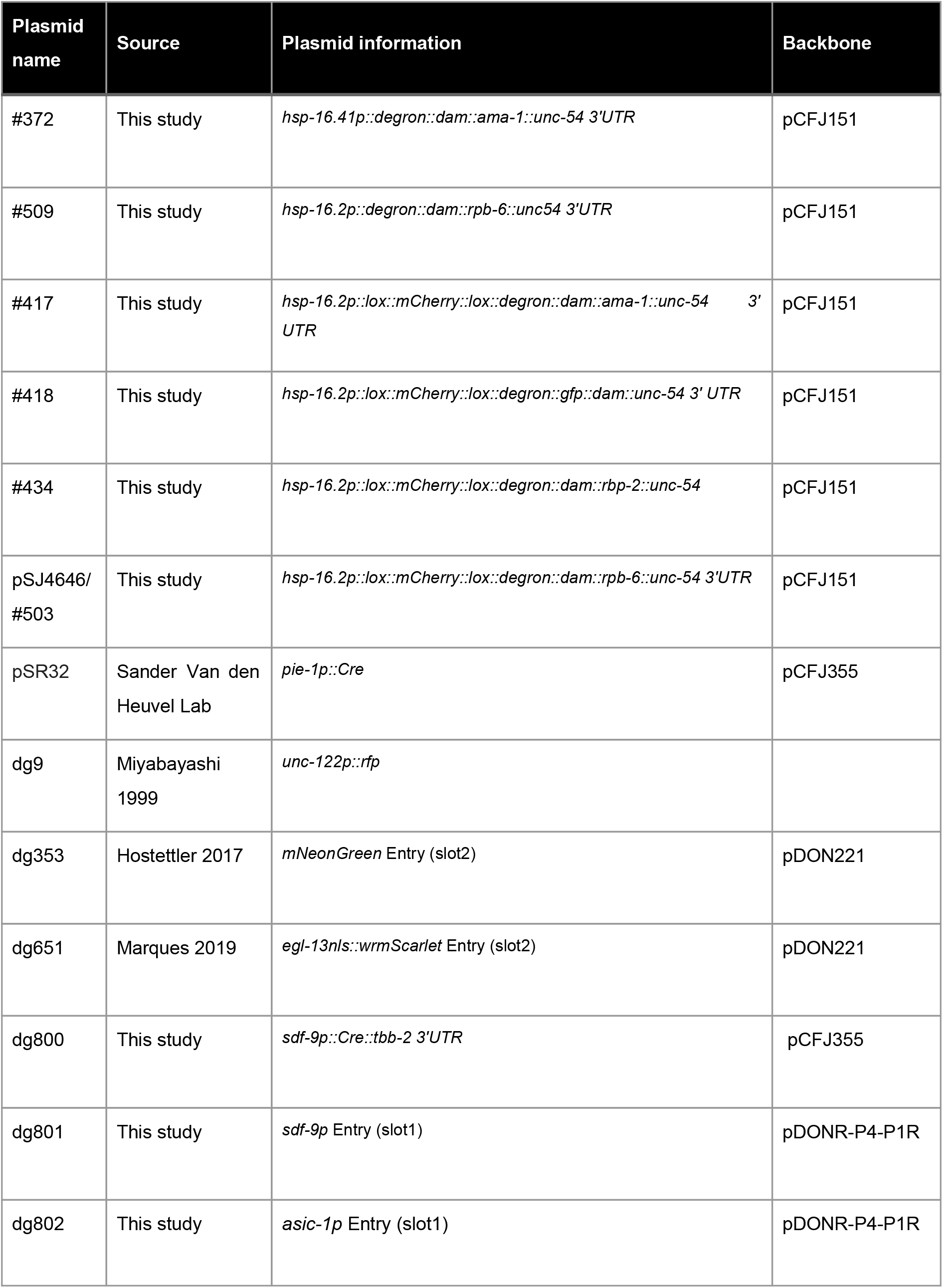
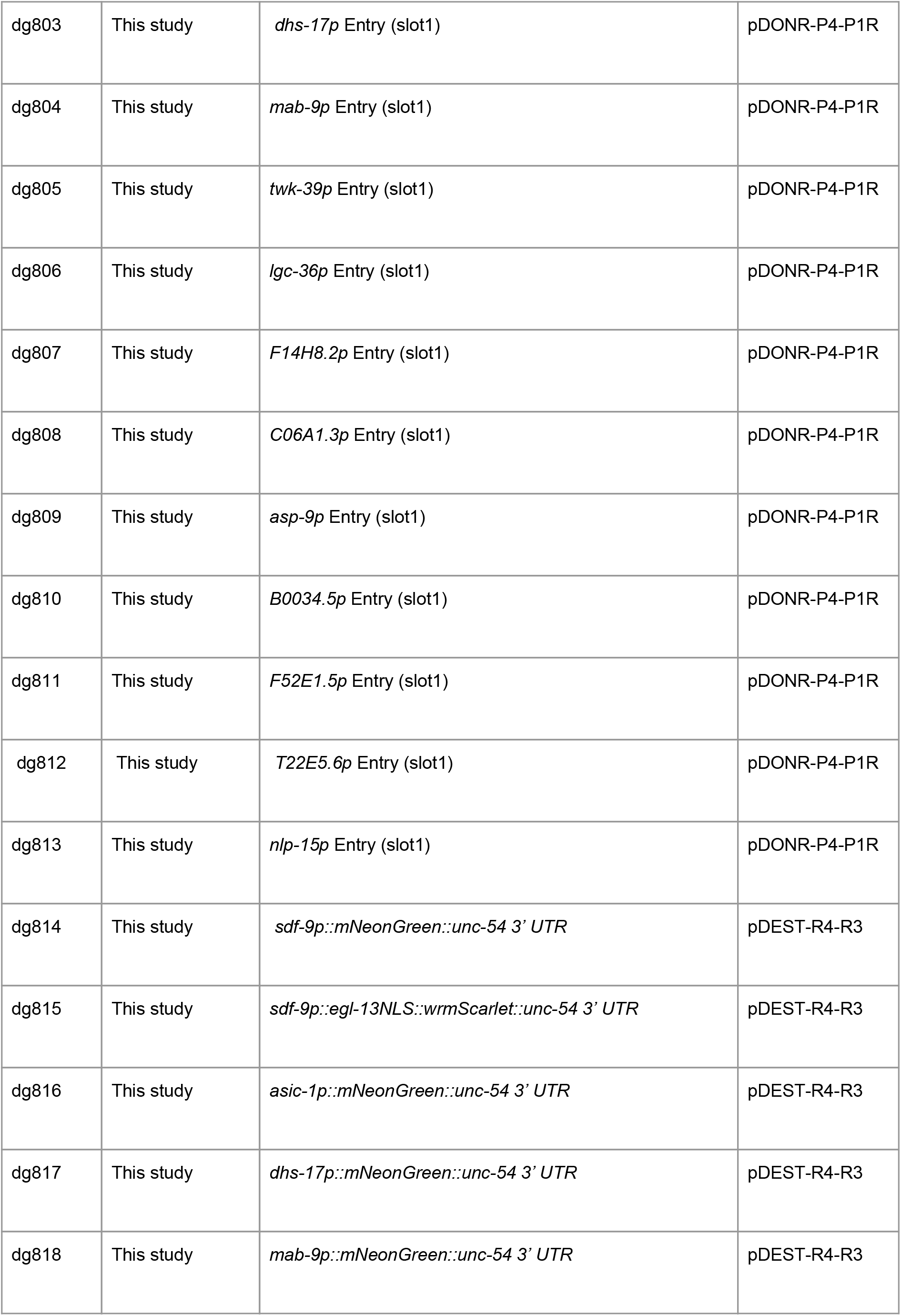
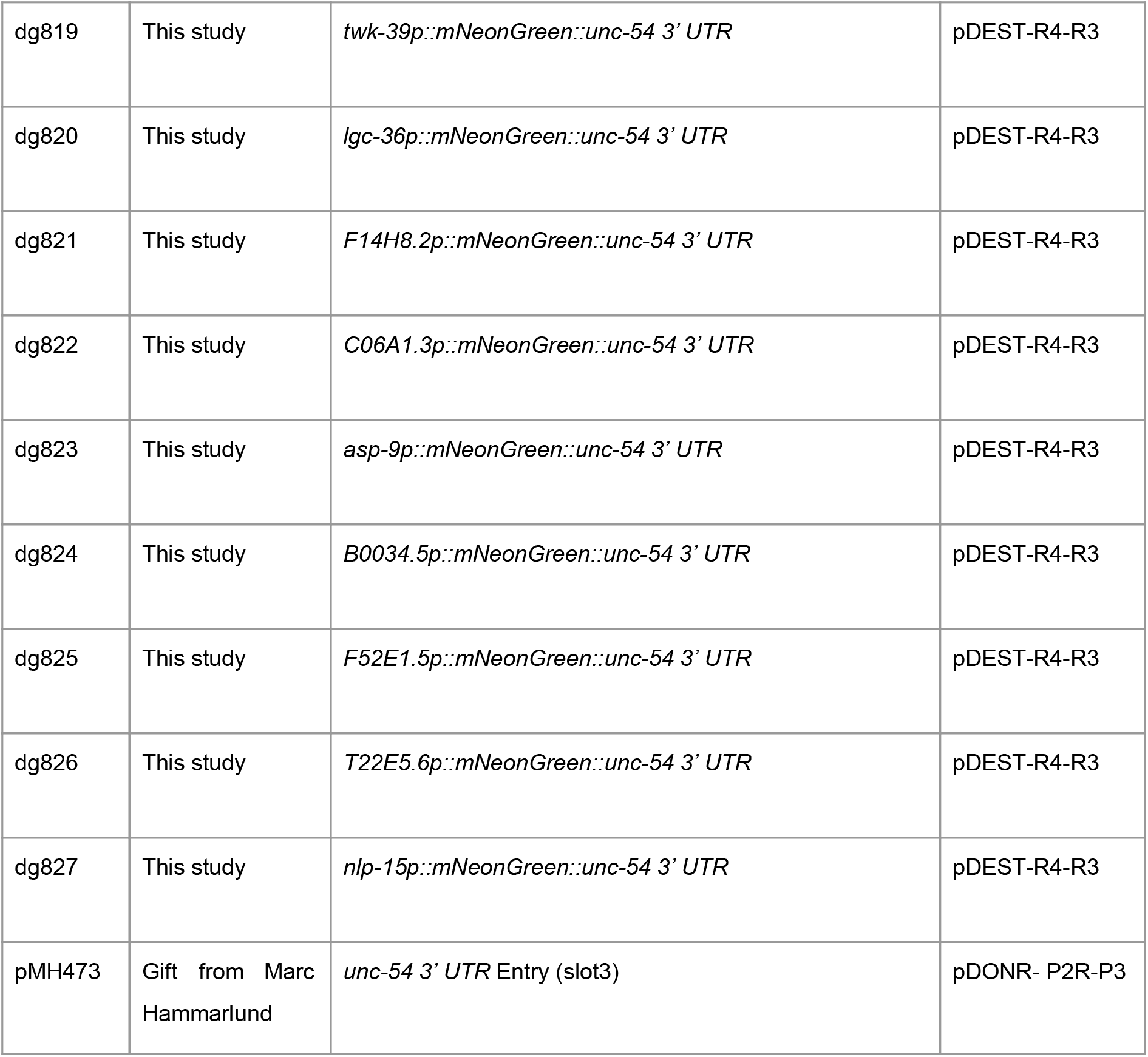
Plasmid collection

**Table S3.**
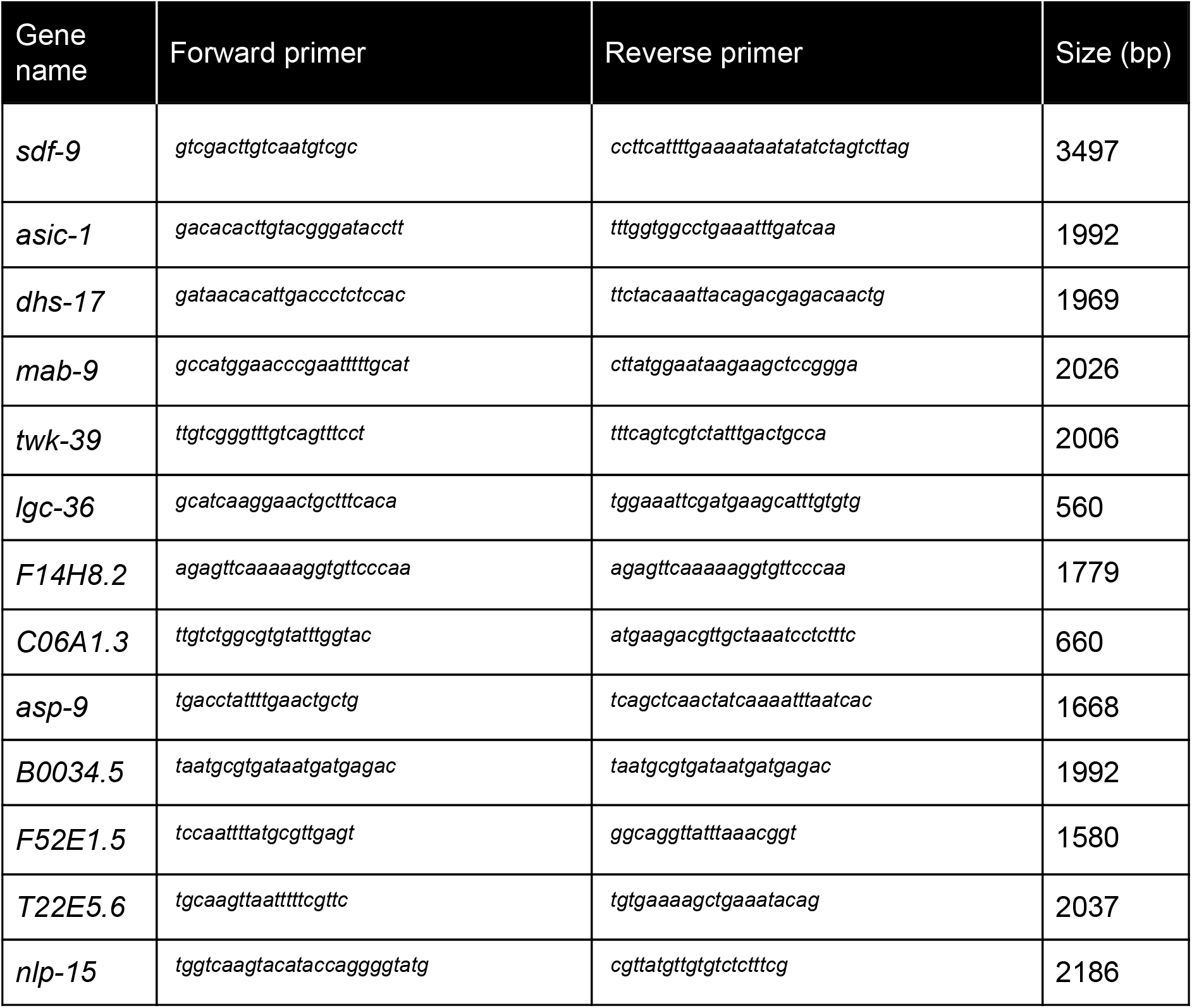
Primers for cloning of promoters used for the transcriptional reporter analysis of XXX cell-transcribed candidate genes. All primers start with Gateway cloning sequences, followed with the indicated sequence. (forward: ggggacaactttgtatagaaaagtt; reverse: ggggactgcttttttgtacaaacttg)

Table S4 (excel sheet)

Averaged gene-level values for RNA polymerase signal (polii), GATC number and false discovery rate (FDR) for all blastomere libraries.

Table S5 (excel sheet)

Transcribed genes analysis determined by RAPID. Genes for each tissue are listed.

Detected and uniques genes, and Gene ontology attributes are tabulated in different sheets.

Table S6 (excel sheet)

List of categories found in the functional enrichment analysis for genes detected with different methods in the intestine and muscle.

Table S7 (excel sheet)

Comparison between the different methods for cell-type specific transcriptome profiling.

Table S8 (excel sheet)

Sequencing depth for the RAPID libraries.

